# CD44-DT regulates p21-dependent senescence via interacting with AR-PCNA complex, revealing a therapeutic vulnerability to chimeric ncRNA-mediated AR degradation in prostate cancer

**DOI:** 10.64898/2026.11.17.767372

**Authors:** Forough Hakiminia, Julia Kallenbach, Katrin Schindler, Golnaz Atri Roozbahani, Norman Rahnis, Aria Baniahmad, Malek Hossein Asadi

## Abstract

Aberrant androgen receptor (AR) signaling driven by androgens plays a critical role in prostate cancer (PCa) progression. Interestingly, bipolar androgen therapy (BAT), a novel therapeutic strategy with cycles between near-castrate and supraphysiological androgen levels (SAL), suppresses PCa growth. SAL induces cellular senescence in various PCa models mediated by the AR, activation of AKT signaling and hypophosphorylation of pRb. Here, we show that SAL upregulates the lncRNA CD44-DT, the CD44 divergent transcript, in PCa cells linking CD44-DT to AR signaling. Knockdown and overexpression datasets suggest that CD44-DT supresses SAL-induced senescence as a negative feed-back loop. Transcriptome, ATAC-seq, and ChIP-seq analyses of PCa cells reveal that AR is recruited to the first intron of CD44 gene locus. Transcriptome analyses of the knockdown suggests that members of AP-1 family act in concert with AR as an upstream activator of CD44-DT. Accordingly, inhibition of the AP-1 complex by a small molecule inhibitor results in downregulation of CD44-DT expression, enhanced SAL-induced senescence and reduced proliferation. Mechanistically, CD44-DT physically interacts with PCNA and AR. This interaction facilitates p21 tumor suppressor degradation, permits cyclin-CDK activation, retinoblastoma (Rb) hyperphosphorylation-activation of the AKT-p70S6K pathway. These findings suggest that CD44-DT is part of AR signaling and acts as a novel onco-lncRNA that is upregulated by AR. To validate the role of AR, we engineered a chimeric ncRNA capable degrading the AR as a novel type of RNA-PROTAC. Accordingly, this RNA-based AR-degrader downregulates CD44-DT confirming AR regulation of CD44-DT. Interestingly, the RNA-PROTAC potently induces apoptosis. Thus, the data suggest that CD44-DT is a novel direct AR target gene that modulates PCNA and p21 protein stability identifying a novel negative feed-back loop at the crossroad between proliferation and senescence indicating CD44-DT as a potential therapeutic target for PCa.

## Introduction

Prostate cancer (PCa) is the most frequently diagnosed cancer among men and is the second leading cause of cancer-related death from male cancers in Western European countries [1, 2]. Dysregulation and abnormal activity of androgen receptor (AR) signaling driven by androgens regulate PCa growth [3]. Therefore, AR targeting with androgen deprivation therapy (ADT), mostly combined with AR antagonists, is a main hormone-based therapeutic approach to inhibit AR activity and consequently restrict PCa growth [4]. Unfortunately, patients with PCa often exhibit resistant to these therapies, and tumors develop to castration-resistant PCa (CRPC) where cancer cells continue to proliferate despite the lack or low level of androgens. However, the AR remains mostly a key driver also for CRPC growth [5].

Strikingly, cyclical supraphysiologic androgen levels (SAL) in androgen deprivation background termed bipolar androgen therapy (BAT) is currently evaluated in phase II clinical trials, including TRANSFORMER, RESTORE and COMBAT, for patients with metastatic CRPC [6–8]. Previous studies have shown that SAL induces cellular senescence, a stable cell cycle arrest, that leads to inhibition of PCa growth. Interestingly, lncRNAs emerge as one driver of SAL-mediated regulation, however, the exact mechanisms are unclear. SAL-mediated senescence was shown to be mediated via signaling pathways such as p15 ^INK4b^/p16^INK4a^/pRb, AKT, and RBL2-DREAM complex signaling pathways [4, 9–12].

In CRPC, transition from an AR-dependent state to an AR-negative state leads to resistance to AR targeted therapies. A recent report identified a subtype of CRPC that develops from AR-dependent to AR-negative stem cell-like (SCL) subtype termed CRPC-SCL. Of note, CRPC-SCL is driven by activator protein-1 (AP-1) family members. CRPC-SCL express high levels of the cancer stem cell marker CD44 [13]. CD44 is a major transmembrane glycoprotein that, through interaction with hyaluronic acid, activates PI3K-AKT pathway, thereby promoting cell proliferation [14, 15].

LncRNAs are defined as non-coding RNAs longer than 200 nucleotides that lack protein-coding ability but play key roles in various cellular processes [16]. The gene encoding the CD44 divergent transcript, CD44-DT, is coding for a long non-coding RNA (lncRNA) arranged in a head-to-head orientation with the CD44 gene separated by a 1.1 kb intergenic promoter region. Bidirectional promoters (BDPs) are evolutionarily conserved and responsible for transcription of around 11% of human genes [17]. A subset of BDPs regulates the transcription of a non-coding and a protein-coding genes located on opposite DNA strands. Like the majority of bidirectional promoters, this promoter lacks a TATA box but contains transcription factor binding sites that are frequently observed in bidirectional promoters [18]. Of note, divergently expressed lncRNAs regulate the transcription of nearby protein-coding genes in pluripotent stem cells [17, 19]. Thus, to obtain insights into molecular pathways of therapies for patients with advanced PCa depends on deciphering the exact mechanisms by which coding and non-coding genes drive castration resistance and tumor plasticity.

Here, we observed that the expression of both CD44-DT and CD44 are upregulated in low-grade PCa. CD44-DT expression is positively correlated with CD44 across all analyzed cancer types retrieved from TCGA indicating a common driver for their expression. Interestingly, functional analyses revealed that CD44-DT positively regulates the expression of its neighboring gene CD44. In addition, we identified CD44-DT as a novel SAL-upregulated lncRNA shown in SAL-treated PCa cell lines and C4-2 xenografted mouse model treated with SAL. Moreover, CD44-DT suppresses SAL-induced senescence in 2D adherent cell culture and 3D spheroid models of PCa. Mechanistically, CD44-DT modulates AKT-p70S6K and E2F1-pRb pathways.

Integration of transcriptome analysis with hallmark pathway genes in PRAD and AR ChIP-seq datasets showed a significant association between the expression of CD44-DT and AR-regulated hallmark pathways. Mechanistically, ATAC-seq and AR ChIP-seq datasets analyses suggest that SAL induces CD44-DT upregulation by increasing chromatin accessibility within the first intron of the CD44 locus, thereby enhancing AR recruitment to this regulatory region. Simultaneously, FOSL2, a member of the AP-1 transcription factor family, is also upregulated by SAL and AR recruited to the nearby regulatory genomic region. Functionally we revealed that AP-1 markedly induces CD44-DT transcription in the both AR-positive cell lines, C4-2 and LNCaP, but not in the AR-negative PC3 cell line, suggesting that AP-1-mediated upstream regulation of CD44-DT expression is dependent on AR signaling.

At protein level, the lncRNA CD44-DT interacts with proliferating cell nuclear antigen (PCNA)-AR complex and leads to its stabilization. PCNA stabilization promotes p21 degradation in nuclei of PCa cells, and subsequently activates Cyclin-CDKs, which results in Rb hyperphosphorylation. Thus, it is appear targeting AR and subsequently reducing the AR target genes as well as SAL-induced CD44-DT upregulation could be a potential strategy for therapy against CRPC. So, we designed a novel chimeric ncRNA that acts as a PROTAC to degrade AR and consequently disrupts the AR signaling, which leads to suppression of PCa growth via inducing apoptosis.

Altogether, our findings identify CD44-DT as a novel onco-lncRNA that promotes PCa growth and restricts cellular senescence in an androgen-dependent manner. We also created an innovative chimeric ncRNA platform that acts as a RNA-based PROTAC to facilitate AR degradation, thereby suppressing PCa survival and growth.

## Materials and Methods

### Cell culture and treatments

The human castration-sensitive PCa cell line LNCaP, and the castration-resistant C4-2 cell line were cultured in RPMI and DMEM (Gibco), respectively. The AR-negative and castration-resistant PC3 cell line was cultured in RPMI supplemented with 5% fetal bovine serum (Gibco) at 37 °C in a humidified incubator with 5% CO2. Cells were treated with 1 nM R1881 (Merck; Germany; R0908) used as SAL [9], 30 µM T5224 (AP-1 inhibitor, (MedChemExpress; HY-12270), or with 0.1 % DMSO (ROTH; Germany; 4720.1) used as a solvent control.

### Knockdown and overexpression of CD44-DT in PCa cell lines

To investigate the role of CD44-DT in PCa cells, siRNA-mediated knockdown, siCD44-DT, and vector-based overexpression, oeCD44-DT, experiments were performed. Briefly, knockdown experiments were performed using Human SMART Pool siRNAs directed against the CD44-DT transcript (siCD44-DT, Dharmacon, R-189406-01-0005), or scrambled siRNA as a negative control (siControl, Dharmacon, D-001810-01-05) at a final concentration 25 nM. PCa cells were transiently transfected with siCD44-DT or siControl employing DharmaFECT reagent 3 Transfection Reagent (Horizon Discovery, Cambridge, UK) according to the manufacturer’s instructions. The efficiency of CD44-DT knockdown was determined using qRT-PCR. CD44-DT overexpression was performed by inserting its full length cDNA (purchased from Eurofins Genomics, Ebersberg, Germany) into the pCDH-CMV expression vector using jetPRIME transfection reagent from Polyplus (Sartorius, Göttingen, Germany) according to the manufacturer’s guidelines. Empty vector (EV), the pCDH-CMV vector without CD44-DT, was used as negative control.

### Creation of a chimeric ncRNA as an AR PROTAC

Our design strategy for the PROTAC chimeric ncRNA (cncRNA) was constructed of the functional protein-interacting domain from the lncRNA HOTAIR interacting with DZIP3 and the lncRNA SLNCR1 sequence interacting with the AR. In order to gain insights into sequence-structure-function relationships of these transcripts, we analyzed their sequences according to literature to use their functional domain-related sequences [20–23]. Then, we selected the E3 ubiquitin ligase DZIP3-binding domain of HOTAIR, HOTAIR^948-1087^, and the AR-targeting fragment from SLNCR1 (SLNCR1^597-637^) which were linked via a short sequence linker to generate a vector expressing a fusion cncRNA. The numbers indicate nucleotide positions. HOTAIR^948-1087^, SLNCR1^597-637^ and cncRNA constructs were synthesized (purchased from Eurofins Genomics, Ebersberg, Germany) and inserted into the pCDH-CMV vector and subsequently transfected into cells using jetPRIME transfection reagent (Polyplus, Sartorius, Göttingen, Germany) in accordance with the manufacturer’s guidelines. HOTAIR^948-1087^, SLNCR1^597-637^ and the empty vector were used as negative controls. The sequences of these constructs are listed in Table S1.

### In vitro transcription

In vitro transcription (IVT) was performed in accordance with manufacturer’s guidelines. Briefly, linearized DNA containing the T7 promoter upstream of target genes was used as template and IVT was performed using HiScribe T7 High Yield RNA Synthesis Kit (Catalog No. E2040).

### RNA pull down assay

Biotin-UTP (Biotin16-UTP, NU-821-BIO16; Jena Bioscience) was used for biotin-labelled full length of CD44-DT was in vitro transcribed as above described and then used for RNA pull-down assay. Next, cell harvested and nuclear lysate protein was prepared and incubated with streptavidin (Novagen, Cat. No. 69203; Merck Millipore), beads and biotin-labeled CD44-DT at 4 °C gently rotation for 2 h. Finally, after five times washing with NT2 buffer containing protases and RNase inhibitors, the eluted proteins were separated on SDS-PAGE, and stained with Coomassie blue and the related bands were analyzed by mass spectroscopy. The biotin-labeled CD44-DT antisense and beads used as negative controls.

### Mass spectroscopy

The bands from the Coomassie stained SDS-PAGE gels were excised and underwent several subsequent steps of dehydration with ACN and rehydration with different aqueous buffer. Next, trypsin was added and the gel pieces were allowed to swell on ice for 30 min, then digested overnight at 37 °C with shaking. After digestion, the peptide content was extracted and dried using a speed vacuum centrifuge (Eppendorf Concentrator Plus, Eppendorf AG, Germany). Peptides were separated using the Evosep One system (Evosep, Odense, Denmark) and were run with a pre-programmed proprietary Evosep gradient of 44 min (30 samples per day) using water and 0.1% formic acid and solvent B acetonitrile and 0.1% formic acid as solvents. The peptides were introduced into the mass spectrometer and the radio frequency ion funnel was set to 30%. For DIA data acquisition, full scan mass spectrometry (MS) spectra with a mass range of 350–1650 m/z were acquired in profile mode in the Orbitrap with a resolution of 120,000 FWHM. For data acquisition and processing of the raw data, Xcalibur 4.4 (Thermo) and Tune version 4.0 were used. DIA raw data were analyzed using the directDIA pipeline in Spectronaut v.21 (Biognosys, Switzerland) with BGS settings besides the following parameters: Protein LFQ method= QUANT 2.0, Proteotypicity Filter = Only proteotypic, Major Group Quantity = Median peptide quantity, Minor Group Quantity = Median precursor quantity, Data Filtering = Qvalue, Normalizing strategy = Local Normalization. The data were searched against a species specific (Homo Sapiens, 20,375 entries, v. 210401) and a contaminants (247 entries) Swissprot database. The candidates and protein report tables were exported from Spectronaut and used for volcano plots generation and principal component analysis (PCA), respectively, using R version 3.4.1 and RStudio server version 1.1.463. Protein groups were considered as significantly enriched in if they displayed a Q value < 0.01 and average log2 ratio > 1.5.

### Co-immunoprecipitation assay

Co-immunoprecipitation (Co-IP) assay was performed as previously described [24]. Briefly, specific antibody against AR and normal rabbit IgG, used as a negative control, was bound with magnetic beads. Whole cell lysates were mixed with the antibody-bound beads and incubated at 4 °C for 2h to form protein–protein interactions. Next, five times washing with 1x PBS were performed and beads were resuspended in SDS buffer. The eluted protein complexes were separated on 12 % SDS-PAGE gels and detected by Western blotting. The antibodies list is provided in supplementary Table S2.

### RNA isolation, cDNA synthesis and quantitative RT-PCR (qRT-PCR)

Total RNA was isolated from cells using the RNA-solvent reagent (Omega, USA; R6830) according to the manufacturer’s instructions. Then, first-strand cDNA was synthesized using 1 µg of RNA and the High-Capacity cDNA Reverse Transcription Kit (Applied Biosystems; Lithuania; 4368814). qRT-PCR was performed using the SsoAdvanced Universal SYBR Green Supermix (Bio-Rad, Munich, Germany) with Bio-Rad CFX96 Real-Time PCR Detection System. The obtained data were normalized to GAPDH and HPRT gene expression and relative gene expression was calculated using the 2^−ΔΔCt^ method. Primer sequences are listed in Table S3.

### Senescence-associated beta-galactosidase (SA-β-Gal) staining

Senescent cells were detected by SA-β-Gal staining, as previously described [9]. Briefly, cells were fixed with 1% glutaraldehyde and subsequently incubated at 37 °C in staining solution containing 40 mM citric acid/sodium phosphate, 5 mM potassium ferricyanide, 5 mM potassium ferrocyanide, 2 mM MgCl₂, 2 mM NaCl, and 50 mg/ml X-gal. Positive stained cells were counted under light microscopy (Carl Zeiss AG, Oberkochen, Germany).

### Growth assays

Cell growth was analyzed by crystal violet staining as previously described [10]. Briefly, cells were fixed with 1% glutaraldehyde and stained with 0.1% crystal violet. The excess stain was washed with Sörensen’s solution containing 0.9% w/v trisodium citrate, 2% HCl, and 40% ethanol in water. Absorbance was measured at 590 nm.

### Protein extraction and Western blotting

Cell pellets were lysed in NETN buffer containing 20 mM Tris-HCl (pH 8.0), 100 mM NaCl, 1 mM EDTA, 1% Tergitol (NP-40), including the phosphatase inhibitors 50 mM NaF, 100 µM Na₃VO₄, and 10 mM β-glycerophosphate. A BCA assay (Thermo Fisher Scientific) was performed to determine protein concentration. Thirty micrograms of isolated proteins were separated by SDS-PAGE and transferred to a membrane. Immunodetection was performed using an Image Quant™ LAS 4000 system (GE Healthcare Bio-Sciences AB, Germany). Band intensities were quantified relative to housekeeping proteins using LabImage D1 software. Primary antibodies are listed in Table S2.

### Tumor spheroid generation, SA-β-Gal, and immunofluorescent staining

Three-dimensional (3D) spheroids were generated from PCa cells as previously described [25]. Briefly, one thousand cells per well were seeded in 96-well ultra-low attachment plates (PerkinElmer) and centrifuged shortly at 300 rpm for three times to facilitate spheroid formation and then incubated at 37 °C in a humidified atmosphere containing 5% CO₂. Next, spheroids were treated for six days with DMSO, SAL (1nM R1881), or the AP1 inhibitor T5224. Spheroids were subsequently fixed with 4% PFA and stained with SA-β-Gal staining solution as described above. The following day, they were washed with PBS and paraffin-embedded. Dehydration was performed using Xylol and a graded ethanol series. Heat-induced antigen retrieval was performed in citrate buffer and followed by permeabilization using 0.2% Triton X100. For Ki-67 detection, spheroid sections were blocked in 5% normal goat serum (NGS) (Biozol; Germany; ENG9010-10) and then incubated with the primary antibody overnight at 4 °C. Then, the spheroids were washed with PBS and incubated with the secondary antibody for 1 hour at room temperature (RT). To discriminate the localization of protein signals, spheroids were then stained with DAPI (Life Technologies; USA; H3569) for 10 min and sections were mounted with Fluoromount-G® (SouthernBiotech; USA; 0100–01) and visualized under a fluorescence microscope (Keyence, BZ-X800 Analyzer). We also performed immunofluorescent staining to identify the localization of proteins in cancer cells as described above with a little difference as visualized under a confocal laser-scanning microscope (Carl Zeiss LSM 880). Antibodies used in this study are listed in supplementary Table S2.

### Mouse xenograft experiment

Mouse experiments were approved by Thüringer Landesamt für Lebensmittelsicherheit und Verbraucherschutz, Germany (Reg.-Nr.: UKJ-23–013). C4-2 xenografts were generated by subcutaneous injection of an equal mixture of C4-2 cells with Matrigel (CORNING; USA; 356231) into both flanks of intact (non-castrated) male athymic nude mice (8-week-old, Janvier Labs, France) [10, 24].

### Chromatin immunoprecipitation sequencing (ChIP-Seq) and data analysis

Chromatin immunoprecipitation (ChIP) was performed using the iDeal ChIP-Seq Kit (Diagenode, Cat.-Nr.: C01010055) according to the manufacturer’s protocol as previous described [26]. Briefly, cells were cross-linked using ChIP Cross-Link Gold (Diagenode). Cells were subsequently fixed by 1% formaldehyde and quenched with glycine. Chromatin was then sheared using a Bioruptor Pico to generate 150-900 bp fragments. Immunoprecipitation was performed using protein A-conjugated beads coupled to a ChIP-grade AR antibody (Cell Signaling, Cat.-Nr: 5153). The isolated DNA was used to generate ChIP-seq libraries using the TruSeq ChIP-Seq Kit and sequenced by macrogen (Seoul, South Korea). FastQC (v0.12.1) and Trim Galore (v0.6.10) were employed to evaluate FASTQ quality and perform read trimming, respectively. After aligning reads to the hg38 reference genome using Bowtie2 (v2.3.5.1), duplicate reads were eliminated. ChIP-seq peaks were identified using MACS14 (v1.4.2) with a p-value cutoff of 1 × 10⁻⁵. For downstream analysis, a reference peak set was generated by merging the detected peaks using the TCseq (v1.30.0) R package. We also analyzed alterations in chromatin architecture following DHT treatment in LNCaP cells via transposase-accessible chromatin (ATAC-seq) and ChIP-seq datasets under accession numbers GSE247596 and GSE232554 [27].

### RNA-seq and data analysis

RNA-seq was performed to investigate the impact of CD44-DT silencing with or without SAL on the transcriptome of C4-2 cells. After treating cells, RNA extraction was performed as described in the protocol above and sent for sequencing to macrogen. First, raw RNA-seq reads were quality-controlled and trimmed prior to transcript quantification using the R package *tximport,* and differential gene expression analysis was performed in R using the *DESeq2* package. Gene expression data obtained from TCGA (The Cancer Genome Atlas) and RNA-seq data from patients who received BAT under accession number GSE229555 were analyzed using R version 4.5.1.

### Ubiquitination assay

To confirm the protein ubiquitination, cells were treated with the MG132 (Selleckchem, S2619), a proteasome inhibitor, that leads to accumulation of ubiquitinated proteins through inhibiting proteasomal degradation. Six hours before cell harvesting, the transfected cells were treated with MG-132 at 15 µM. Then, the harvest cells were subjected to protein extraction and western blotting as described previously.

### Molecular docking

The human PCNA were retrieved from the UniProt Knowledgebase (UniProtKB) under the accession number P12004 and its crystal structure was also obtained from the Protein Data Bank (PDB) under the PDB ID 9N3L. The protein structure was obtained from AlphaFold protein structure database (https://alphafold.ebi.ac.uk/). PCNA and CD44-DT molecular docking was performed using the HADDOCK 2.4 web server (https://wenmr.science.uu.nl/had-dock2.4/) [28, 29]. Further structural analysis and visualization were performed by the Chime-raX program [30].

### Statistical analysis

To statistically analyze the obtained data, we employed GraphPad Prism 10.0 (RRID: SCR-000306). A two-tailed unpaired t-test was used to compare two groups, while multiple group comparisons were performed using two-way ANOVA. All statistical data are presented as the mean values ± standard deviation (SD) or standard error of the mean (SEM).

## Results

### CD44-DT is a novel androgen-induced lncRNA that associates with oncogenic hallmark pathways in PRAD

Divergent lncRNAs are a class of lncRNAs transcribed from BDPs comprising approximately 20% of all lncRNAs in mammalian genomes. Divergent lncRNAs in pluripotent stem cells have been identified to regulate the transcription of their protein-coding partners, suggesting a fine-tuning of gene expression [19]. However, the regulatory mechanisms of divergent transcripts remain poorly understood. We hypothesized that the expression of CD44 and its divergent transcript, CD44-DT, co-regulated. To address that hypothesis, co-expression of CD44-DT and CD44 was analyzed in 32 tumor types in pan-cancer datasets. Separately, correlation analyses were performed in prostate adenocarcinoma datasets (PRAD) and the ENCORI database. In the ENCORI datasets, the identification number of CD44-DT is recorded as AL356215.1 that is verified by an unique Ensmbl gene ID, ENSG00000255521. Interestingly, a markedly positive correlation between CD44-DT and CD44 expression at RNA levels in all analyzed patient cancer types was retrieved (Fig. 1a) including PCa (Fig. 1b). Correlation analysis of TCGA PRAD dataset further revealed a significant association between CD44-DT expression and hallmark signaling pathways implicated in PCa growth, progression, and therapy resistance, such as NF-κB, p53, PI3K-AKT-mTOR, Wnt/β-catenin, Notch, and androgen response signaling (Fig. 1c). We further examined the association between CD44-DT expression and stemness/senescence-associated gene expression in PRAD datasets. Notably, CD44-DT is significantly higher expressed along with upregulation of stemness-associated genes. Notably, CD44-DT expression also positively correlates with the expression of senescence-associated genes (Fig. 1d).

**Fig. 1:**
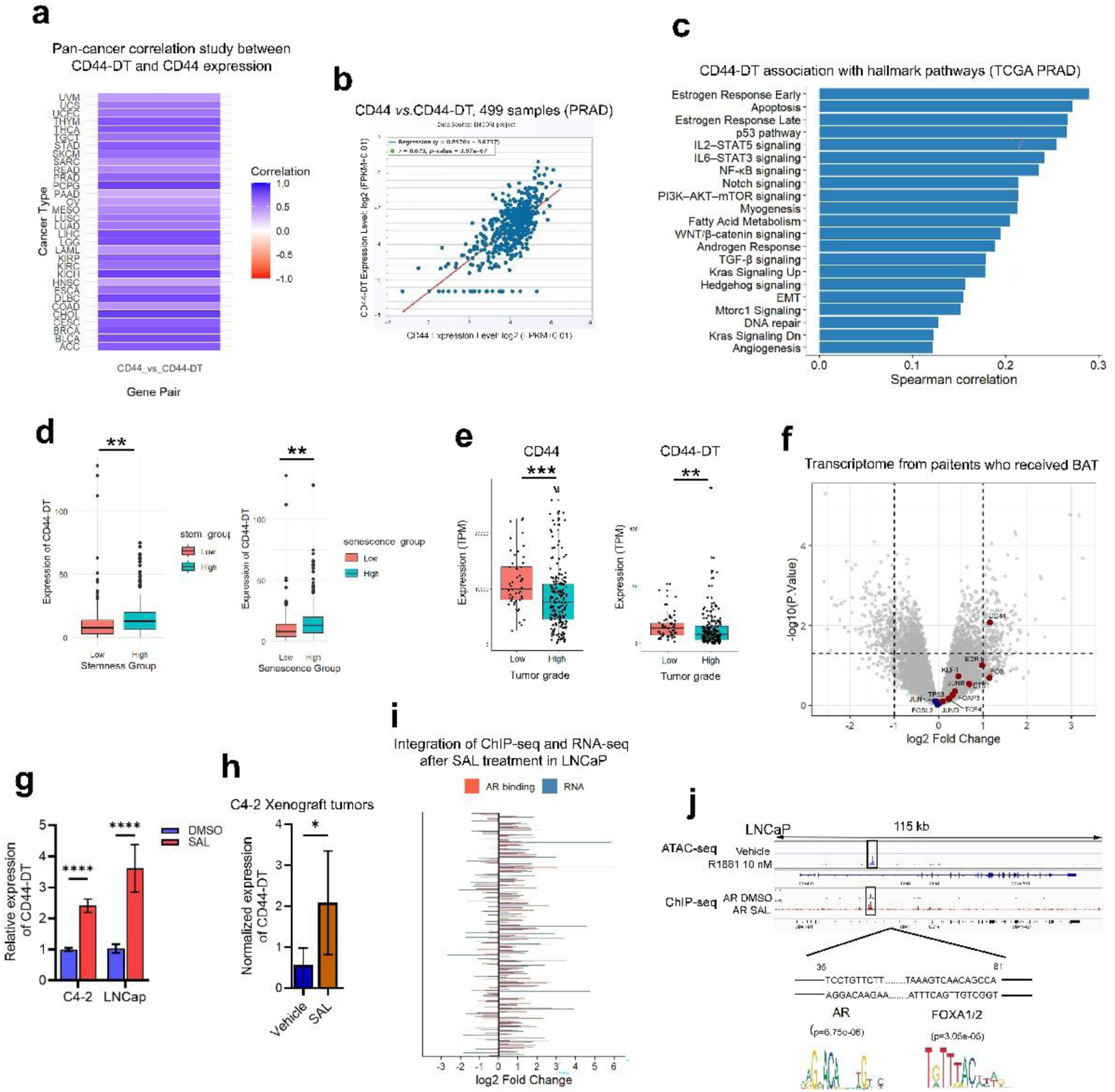
CD44-DT expression associated with oncogenic hallmarks pathways in PCa. **a)** TCGA data analyze showing the positive correlation between the expression of CD44-DT and CD44 in 32 cancer types **b)** including PCa (R = 0.673, p = 3.97e-67). **c)** Significant oncogenic hallmark gene sets identified by Gene Set Enrichment Analysis (GSEA) related to CD44-DT expression in PRAD cohort. **d)** The significant positive correlation between the expression of CD44-DT and stemness/senescence-associated genes in PRAD datasets. **e)** Boxplots view for PRAD datasets analysis showing the significant upregulation of CD44 and CD44-DT in low-grade PCa compared to high-grade ones. **f)** Volcano plot of RNA-seq analysis reveals that CD44 is upregulated in PCa patients who received BAT. **g)** CD44-DT upregulation in SAL-treated PCa cells and **h)** in the C4-2 xenograft mice model treated with SAL (n= 4) compared to vehicle (n = 5) was detected using qRT-PCR. Data represent the mean± SEM. **i)** A tornado diagram illustrates the integration analysis of AR ChIP-seq and transcriptome data from SAL-treated LNCaP cells highlighting androgen-responsive genes within hallmark pathways. **j)** Genomic tracks showing ATAC-seq accessible peaks and ChIP-seq signal at CD44 genomic loci in response to SAL compared to DMSO visualized with IGV software in LNCaP cells.

The observed correlation between the expression of CD44-DT and stemness-associated genes in PCa prompted us to also investigate its putative association with tumor grades. Paradoxically, PRAD analysis suggests that both CD44-DT and CD44 expression are significantly upregulated in low-grade prostate tumors and their expression are significantly decreased in high-grade tumors (Fig. 1e). This may indicate either a tumor suppressive role of CD44-DT or its importance in early stages of PCa tumorigenesis.

To further investigate the expression of CD44-DT in patient samples who received BAT, the expression of CD44-DT, CD44 and their upstream regulators was analyzed in RNA-seq datasets. The volcano plot of transcriptomes suggests that CD44 is upregulated in patients who received BAT (Fig. 1f). However, the expression of CD44-DT was not detectable in this dataset, likely due to its low expression level as generally, lncRNAs appear to be expressed at lower levels compared to mRNAs [31]. Based on the upregulation of CD44 in patients who received BAT and the positive correlation between CD44-DT and CD44 expression likely due to the shared BDP, we hypothesized that CD44-DT expression, similar to CD44, is induced by androgen in PCa. Therefore, to confirm whether CD44-DT is categorized as an androgen-responsive lncRNA, its expression was measured following androgen treatment at SAL in PCa cell lines and C4-2 xenografted mice model treated with SAL in vivo. In line with the hypothesis, gene expression analysis revealed that CD44-DT was significantly upregulated in the SAL-treated PCa cell lines, LNCaP and C4-2, as well as in the C4-2 xenograft mice model treated with SAL compared with control ones treated with DMSO or vehicle, respectively (Fig. 1g and 1h). This suggests that CD44-DT is part of androgen signaling.

To obtain more insights into the transcriptional regulation, integrative analysis of ChIP-seq and RNA-seq datasets from SAL-treated LNCaP cells was performed (Fig. 1i). Tornado plot supports a co-expression of CD44 and CD44-DT being within AR signaling in PCa cells. The analysis of ATAC-seq data from LNCaP cells (GSE247593) identified a potential cis-acting regulatory element within the first intron of CD44 that becomes more accessible upon SAL treatment, suggesting that this genomic region opens up in response to SAL and likely triggers CD44-DT transcription. In line with this, our AR ChIP-seq data also showed a peak at the same genomic region by SAL treatment suggesting the recruitment of AR to this region. Motif binding site prediction analysis with JASPAR and FIMO identified high-confidence adjacent binding sites for AR and FOXA1/2 at this region suggesting that AR binds to this region to regulate the expression of CD44-DT (Fig. 1j). FOXA1/2 proteins act as pioneer factors that increase the accessibility for other TFs including AR through chromatin remodeling at enhancer sites [32, 33]. These findings support the mechanistic basis of co-expression of CD44 and CD44-DT at chromatin level.

### Crosstalk between AP-1 and AR signaling as upstream regulators of CD44-DT

The CD44/CD44-DT BDP was already defined above has been suggested to be regulated by upstream transcriptional activators including AP-1 [18, 34] (Fig. 2a). Our above analysis indicates that CD44 is upregulated in patients who received BAT. Also, the expression of CD44-DT is upregulated by SAL. In addition, RNA-seq analysis of C4-2 and LNCaP cells treated with SAL indicate that FOS and FOSL2, two members of AP-1 family, are upregulated upon SAL treatment (Fig. 2a). To obtain more insights into the transcriptional regulation, analysis of ATAC-seq and AR ChIP-seq datasets from SAL-treated LNCaP cells suggest the recruitment of AR to nearby FOS and FOSL2 loci (Fig. 2b). Therefore, we hypothesized that SAL-activated AR triggers CD44-DT transcription either directly through AR recruitment to the first intron of CD44 genomic region and additionally in an indirect manner by upregulation of AP-1 family members via AR recruitment to accessible regulatory elements within subunits of AP-1 gene loci. Our ChIP-seq AR confirms the recruitment of AR to members of the AP1 family members (Fig. 2b) indicating that AR upregulates AP1 family members. These data indicate that the upregulation of CD44, CD44-DT by AR is a dual pathway by AR being recruited to AP1 family members as an upstream regulator of CD44-DT and is recruited to the CD44 / CD44-DT locus.

**Fig. 2).**
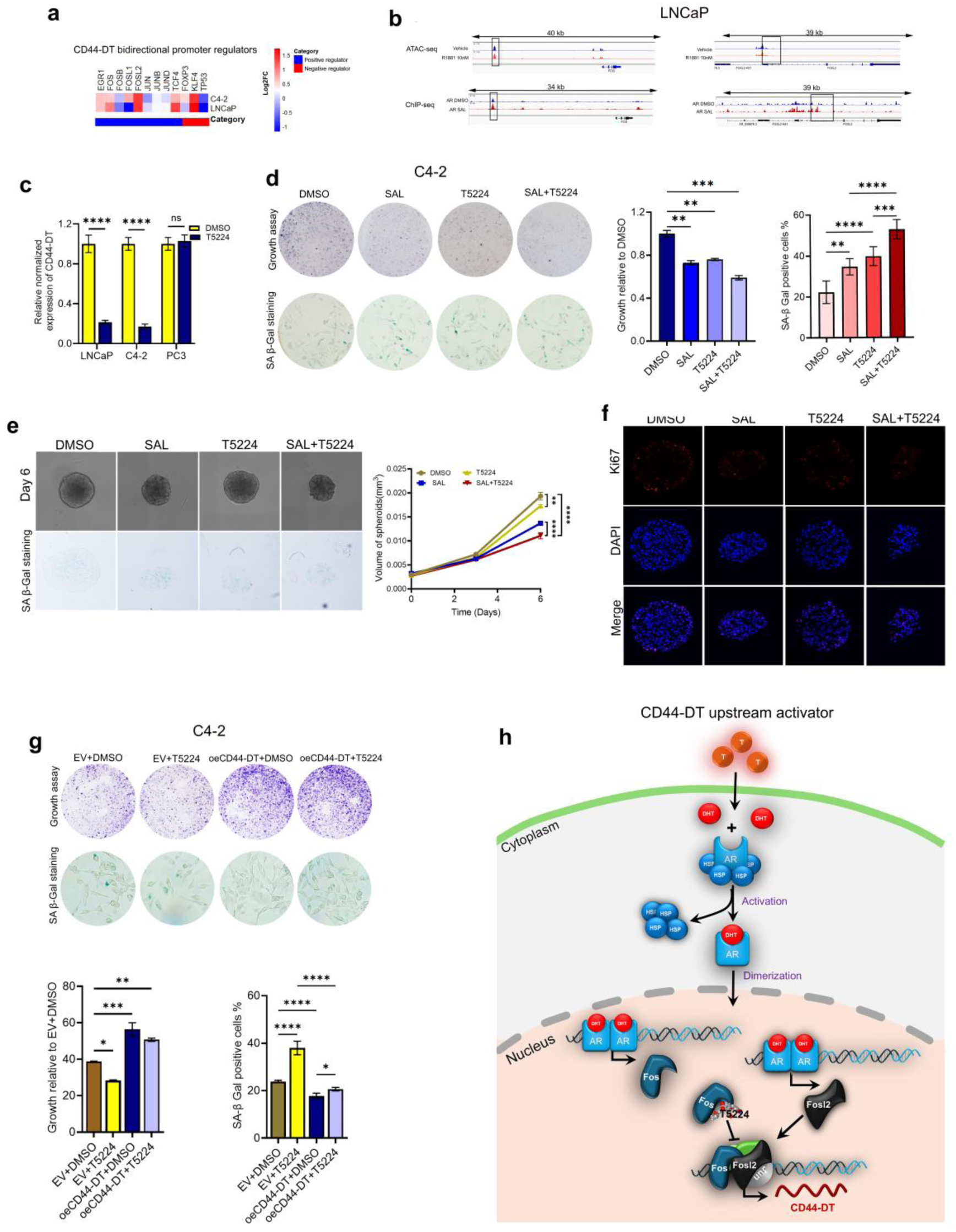
Androgen-responsive AP-1 complex acts as an upstream activator of CD44-DT. a) Heatmap from RNA-seq analysis shows the expression pattern of upstream regulators of CD44-DT in C4-2 and LNCaP cells treated with SAL or DMSO. b) A genome browser view of ATAC-seq accessible peaks and ChIP-seq signal at FOS and FOSL1 loci, two members of AP-1 family, in LNCaP cells following DHT treatment. c) Bar graph shows CD44-DT expression in PCa cell lines following T5224 treatment. d) Crystal violet and SA-β-Gal staining represent growth restriction and senescence induction following T5224 treatment in PCa cells in SAL context. Two-way ANOVA was performed for statistical analysis (*p ≤ 0.05, **p ≤ 0.01, and ***p ≤ 0.001). e) Representative images show the impact of T5224 on C4-2-derived spheroids size. Scale bars correspond to 100 μm. The mean ± SEM values are represented (n = 2, each containing ten technical replicates). Two-way ANOVA was performed for statistical analysis (*p ≤ 0.05, **p ≤ 0.01, ***p ≤ 0.001, and ****p ≤ 0.0001). f) Immunofluorescence staining of C4-2-derived spheroids treated with T5224 shows Ki67 proliferation marker (red), DAPI staining the nucleus (blue), and the merged image. Negative controls are the samples without primary antibody. g) Crystal violet and SA-β-Gal staining represent CD44-DT overexpression rescued T5224-induced growth restriction and cellular senescence in PCa cells. Two-way ANOVA was performed for statistical analysis (*p ≤ 0.05, **p ≤ 0.01, and ***p ≤ 0.001). h) Schematic diagram illustrated the upstream regulation of CD44-DT in response to androgen treatment in PCa cells.

To further support this hypothesis, the AP-1 small molecule inhibitor, T5224, was employed which interferes with the DNA-binding domain of JUN/FOS family members that has reached Phase II clinical trials for rheumatoid arthritis [13, 32]. Treatment of T5224 in LNCaP, C4-2, and the castration-resistant AR-negative PC3 cells revealed that, CD44-DT was markedly downregulated in LNCaP and C4-2 cells by T5224, whereas its expression remained unchanged in PC3 cells. This suggests a cooperation between AR signaling and the AP-1 in regulating CD44-DT expression (Fig. 2c). Interestingly, a growth inhibitory effect of T5224 was observed in all three PCa cells, which was more pronounced in C4-2 cells compared to LNCaP and PC3 cells. Interestingly, T5224 induces cellular senescence in all three PCa cell lines indicating a cellular response to the treatment with T5224 (Fig. 2d, S1a and S1b). Despite the critical role of AP-1 in proliferation of PC3 cells, it does not appear that T5224 regulates CD44-DT transcription in these AR negative cells. This indicates that AR signaling is central to CD44-DT induction and upstream of AP1.

To further validate, the impact of T5224 for the growth of C4-2-derived 3D spheroids and induction of cellular senescence was analyzed. The obtained results suggest that T5224 markedly enhanced SAL-induced senescence and inhibited the volume of C4-2-derived spheroids (Fig. 2e). Ki67 immunofluorescence also confirmed the inhibitory effect of T5224 on C4-2 growth (Fig. 2f). To obtain further insights into this signaling CD44-DT overexpression was used as a rescue experiment for inhibitory effect of T5224. Overexpression of CD44-DT in C4-2 cell in presence of T5224. As expected, CD44-DT rescued both the inhibitory impact of T5224 in cancer cells growth and its inductive effect on cellular senescence (Fig. 2g, p<0.0001).

Taken together, these data support a model in which a novel AR signaling axis as well as a novel feed-back loop are identified. The upstream factor is the SAL-activated AR. AR signaling enhances the expression of CD44-DT and AP1 family members at chromatin level. On one hand CD44-DT counteracts AR activity as one negative feedback loop. The AP1 family members are downstream of AR signaling but upstream of CD44-DT. On the other hand, CD44-DT overexpression ameliorates the effect of the T5224 inhibitor indicating that the senescence induction by T5224 is mediated by downregulation of CD44-DT and suggests that overexpression of CD44-DT loops back to overcome AP1 family member signaling. Schematic diagram represents SAL-activated AR upregulates CD44-DT through activation of AP-1 members (Fig. 2h).

### CD44-DT inhibits SAL-induced cellular senescence and modulates AKT/S6 signaling

To investigate the impact of CD44-DT on growth and SAL-mediated senescence, siRNA-mediated knockdown (KD) was performed in both LNCaP and C4-2 cells. The knockdown efficacy of CD44-DT was determined using qRT-PCR (Fig. S2a) confirming efficient knockdown of CD44-DT expression also in the presence of SAL. Next, RNA-seq and transcriptome analysis was performed with C4-2 cells transfected with siCD44-DT or scrambled siRNA, which were each treated with SAL or DMSO as a control solvent. The gene set enrichment analysis (GSEA) of the transcriptomes suggest that genes repressed by siCD44-DT are enriched in oncogenic programs that are similar to those repressed by SAL including MYC target V1/V2, G2M checkpoint, and E2F targets pathways. In addition, we revisited the well-established role of SAL-mediated cellular senescence using the GSEA obtained from comparing SAL versus DMSO treatment and comparing siCD44-DT versus siControl. Interestingly, siCD44-DT indicates inhibitory impact on SAL-upregulated androgen-response genes (Fig. 3a). This result together with above results suggest a cross-talk between CD44-DT and AR signaling supporting our findings. To confirm whether CD44-DT regulates AR signaling the KD of CD44-DT downregulates the SAL-induced AR target genes, *FKBP5* and *TMPRSS2* (Fig. 3b). This finding suggest that AR induces the expression of CD44-DT, which conversely represses AR target gene expression indicating a negative-feedback loop in which CD44-DT is upregulated by SAL and inhibits SAL-induced expression of AR target genes. Our previous findings established that SAL induces cellular senescence in CSPC and CRPC cells [5, 10, 12, 13, 26] suggested as an important tumor-suppressive mechanism. Consistent with the positive correlation between CD44-DT expression and senescence-associated genes observed in PRAD dataset, the transcriptome analysis revealed that siCD44-DT induces pronounced changes in the expression of senescence pathway genes enhanced by SAL (Fig. 3c). Therefore, we hypothesized that CD44-DT regulates in part SAL-induced cellular senescence. Too address this hypothesis, knockdown (siCD44-DT) and overexpression (oeCD44-DT) experiments were performed with or without SAL treatment in both C4-2 and LNCaP cells. SA-β-Gal and crystal violet staining showed that CD44-DT KD markedly enhanced SAL-induced senescence (Fig. 3d, 3e and S2b) indicating that CD44-DT inhibits SAL-induced cellular senescence.

**Fig. 3).**
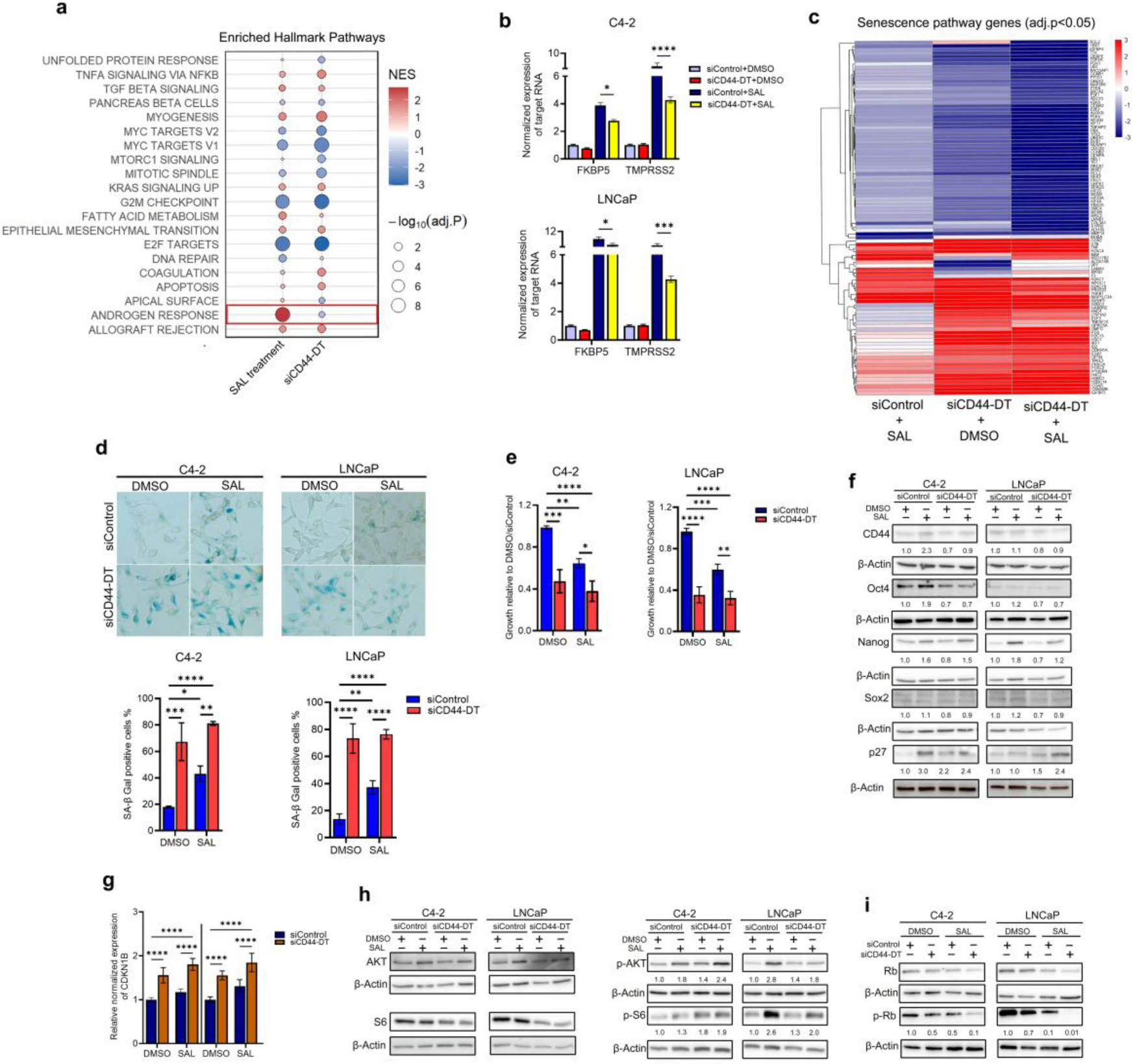
CD44-DT suppresses SAL-induced senescence via regulating the PI3K/AKT/mTOR and pRb-E2F1 signaling pathways. a) GSEA of RNA-seq data from SAL-treated C4-2 cells with or without siCD44-DT based on the hallmark gene set, the red dots indicate significantly enriched pathway regulated by SAL or siCD44-DT in comparison to cells treated with scrambled siRNA plus DMSO. b) Box plots show the impact of siCD44-DT on well-known AR target genes. c) Heatmap showing senescence-associated gene expression from RNA-seq data generating on the Nanostring nCounter platform in SAL-treated C4-2 with or without siCD44-DT versus cells treated with DMSO plus siControl. d and e) The percentage of senescent cells and the ratio growth of PCa cells in presence SAL with or without siCD44-DT compared to cells treated with DMSO plus siControl. Bar graphs are shown as mean ± SD (n = 3). f) Western blot analysis of CD44, core pluripotency transcription factors including, Oct4, Nanog and sox2. Moreover, the expression of CDK inhibitor, p27, in SAL context with or without siCD44-DT at protein level by Western blotting and g) at RNA level by qRT-PCR. Bar graphs are shown as mean ± SD (n = 3). f) Western blot shows the changes in phosphorylation of AR (p-AR), AKT (p-AKT), S6 (p-S6) and Rb (p-Rb) in CD44-DT depleted cells treated with SAL or DMSO. Numbers imply the band intensities that was normalized to β-Actin as an internal loading control. h) Western blotting shows the impact of CD44-DT knockdown on the phosphorylation of Rb in presence of SAL. Number represent the relative band intensity that was normalized to β-Actin (n = 3). Two-way ANOVA was performed for multiple comparisons (n = 3). Data are shown as mean ±SD. *p ≤ 0.05, **p≤0.01, ***p ≤ 0.001, ****p ≤ 0.0001.

In line with the KD results, the overexpression of CD44-DT noticeably reduced cellular senescence (Fig. S2c) and induces growth of C4-2 and LNCaP cells treated with SAL or DMSO (Fig. S2d). This result supports our hypothesis and bioinformatic predictions that CD44-DT regulates cellular senescence and cell proliferation and suggests that CD44-DT acts as an oncogenic lncRNA.

In line with the positive correlation between CD44-DT expression and stemness-associated genes observed in PRAD dataset, the obtained heatmap from RNA-seq data suggests that CD44-DT depletion results in higher expression of stemness-associated genes compared to SAL treatment (Fig. S3a). Therefore, our hypothesis was that CD44-DT controls the expression of stemness regulating genes. To verify the stemness characteristic of CD44-DT, we examined in addition to the expression of its neighboring gene, CD44 that well-known as a cancer stem cells (CSCs) marker, also genes known for core embryonic stem cells (ESCs)-associated TFs including OCT4, NANOG and SOX2 following CD44-DT silencing in absence and presence of SAL. The data indicate that CD44-DT KD represses the SAL-upregulated CD44, Oct4, Nanog and Sox2 in both cell lines (Fig. 3f). CD44-DT Overexpression enhances the expression of Sox2 (Fig. S3c). Thus, these findings suggest that CD44-DT acts also as regulator of stemness-associated pathways in PCa.

The AKT/mTOR pathway has been identified as a pivotal pro-survival mediator of SAL-induced cellular senescence [35]. Interestingly, CD44-DT KD enhances the SAL-induced phosphorylation of AKT and S6 (p70S6K) in castration-resistant C4-2 cells and castration-sensitive LNCaP cells (Fig. 3h). In agreement with knockdown results, CD44-DT overexpression in PCa cells further confirmed its ability to modulate the phosphorylation of AKT and S6 (Fig. S3b).

Cellular senescence occurs when cell cycle arrests through upregulation cell cycle inhibitors, therefore the p27/p21^WAF1/Cip1^-pRb-E2F1 pathway was analyzed. Our results indicate that CD44-DT KD enhances SAL-induced p21^WAF1/Cip1^ and p27 as identified by RNA-seq, Western blotting, and qRT-PCR (Fig. 3c, 3f, 3g and 5e). p21^WAF1/Cip1^ and p27 are cyclin-dependent kinase inhibitors (CKIs) that inhibit phosphorylation of the Retinoblastoma protein (Rb), causing Rb to remain active and inhibit E2F transcription factor, thereby arresting the cell cycle at the G1 checkpoint [36, 37]. Consistently, p-Rb significantly reduced following CD44-DT KD in both LNCaP and C4-2 cells and this reduction is also enhanced by SAL context (Fig. 3i). Accordingly, CD44-DT overexpression induces the phosphorylation of Rb in both PCa cell lines (Fig. S3b) further supporting the data with CD44-DT knockdown. These findings indicate that CD44-DT restricts SAL-induced senescence through regulation of PI3K/AKT/mTOR and p21-pRb-E2F1 signaling pathways in PCa cells.

Altogether, CD44-DT as a novel AR direct target gene, correlates with hallmark pathways promoting PCa growth, tumor progression and therapy resistance including NF-κB, p53, PI3K-AKT-mTOR, and androgen response signaling pathways which also confirmed by RNA-seq analysis (Fig. S4a-d).

### Targeted reduction of the onco-lncRNA CD44-DT inhibits growth of PCa tumor spheroids

Spheroid model better represents the complexity of multicellular tumor architecture, tumor microenvironment and cell-cell interactions in a 3D context compared to 2D adherent culture [10, 11]. Therefore, the impact of CD44-DT silencing with or without SAL treatment was examined for proliferation and cellular senescence in C4-2 and LNCaP cell-derived 3D spheroids. The CD44-DT KD reduced the volume of spheroids derived from C4-2 or LNCaP cells. This indicates that SAL treatment enhances the inhibitory effect of siCD44-DT on the volume of the spheroids (Fig. 4a).

**Fig. 4).**
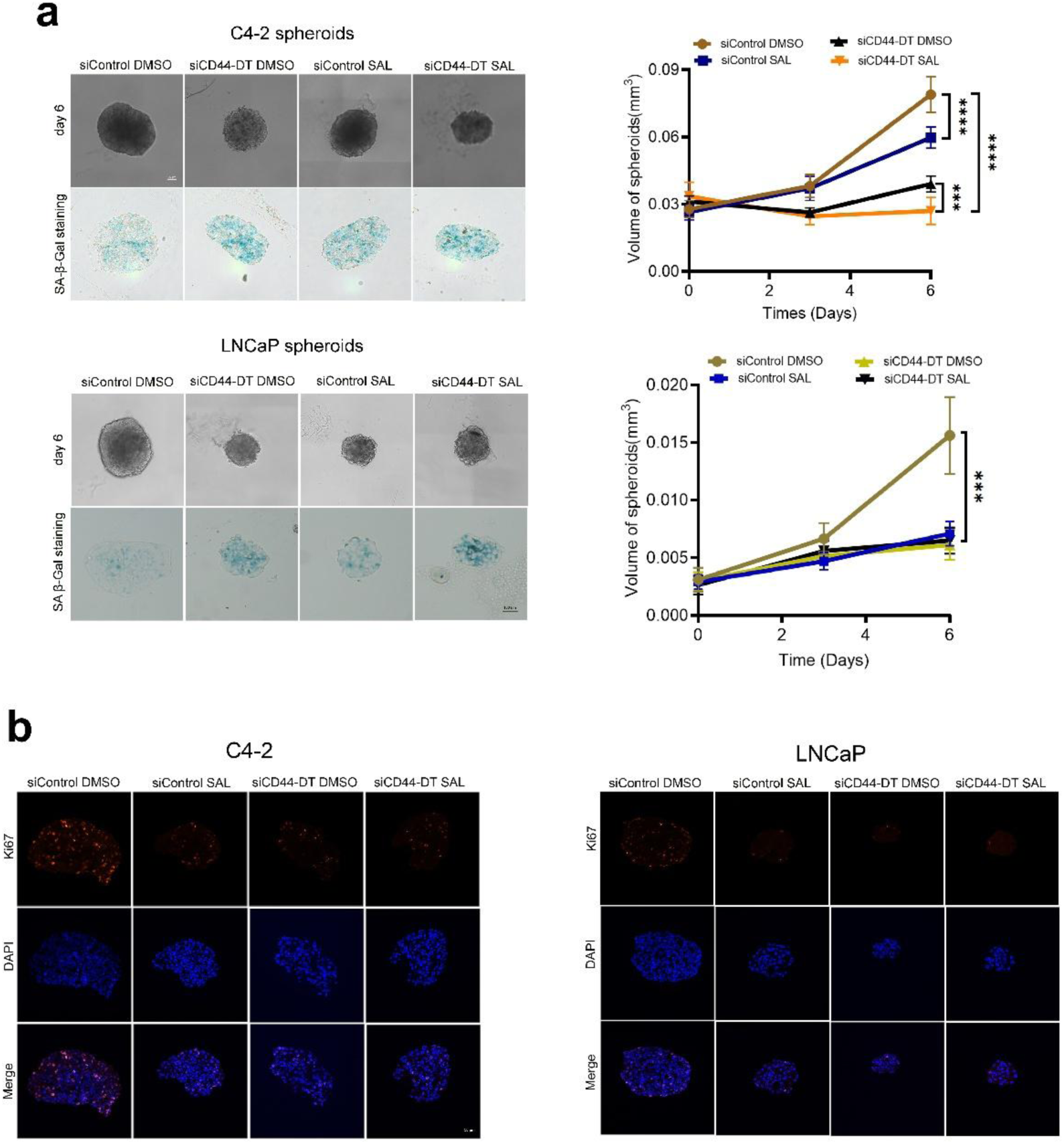
CD44-DT KD reduces the volume of spheroids derived from C4-2 and LNCaP cells. a) Representative images of size and SA-β-Gal staining in CD44-DT depleted spheroids derived from C4-2 and LNCaP in SAL context. Scale bars correspond to 100 μm. The mean ± SEM values are represented (n = 2, each containing ten technical replicates). Two-way ANOVA was performed for statistical analysis (*p ≤ 0.05, **p ≤ 0.01, and ***p ≤ 0.001). b) Immunofluorescence staining shows Ki67 as a proliferation marker (red), DAPI staining the nucleus (blue), and the merged image. Negative controls are the samples without primary antibody. Scale bars correspond to 50 μm.

Similar to the 2D adherent culture, SA-β-Gal staining suggests that the number of senescent cells were increased in knockdown spheroids derived from both cell lines. In line with these findings, immunofluorescence results showed that the expression of Ki67, a well-known proliferation marker, was significantly reduced in siCD44-DT-derived spheroids treated with SAL compared to siControl-derived ones treated with DMSO (Fig. 4b). Thus, these data suggest that CD44-DT inhibits cellular senescence, controls PCa proliferation and acts within AR signaling.

### CD44-DT interacts with PCNA-AR complex and facilities p21 degradation

To obtain further insights into molecular mechanism of CD44-DT in regulating PCa proliferation and senescence, we performed RNA pull-down to identify CD44-DT-interacting proteins (Fig. 5a) by mass spectroscopy (MS-spec). C4-2 cell extracts were incubated with in vitro transcribed biotin labeled CD44-DT coupled beads. Proteins extracted from CD44-DT pull-down assays were subjected to mass-spec and compared with those obtained from antisense RNA control and beads alone as negative controls. Comparative proteomics analysis revealed that several candidate proteins that were specifically detected in the CD44-DT pull-down and not in the negative controls (Fig. 5b). We focused on the proliferating cell nuclear antigen (PCNA) as a key factor of cell proliferation and replication. The interaction of CD44-DT with PCNA was validated by using RNA pull down with subsequent anti-PCNA antibody via Western blotting (Fig. 5c). These data suggest the binding of CD44-DT to PCNA, a key factor of cell proliferation. Based on above results we speculated CD44-DT may interact with AR, and in line with our speculation this interaction verifies by RNA pull down with subsequent anti-PCNA antibody via Western blotting (Fig. 5c). We also hypothesized PCNA and AR may interact with each other and co-immunoprecipitation (Co-IP) assay confirmed our hypothesis. Therefore, our findings showed that CD44-DT physically interacts with PCNA-AR complex (Fig. 5d). To decipher the functional effects of this interaction, we hypothesized CD44-DT may control PCNA protein stability. To address this hypothesis, the level of PCNA was reduced by siCD44-DT and MG132 proteasome inhibition assays suggest CD44-DT might stabilize PCNA (Fig. 5e). Previous reports demonstrated that PCNA binds to p21 and regulates its ubiquitination and degradation [38, 39]. Therefore, we analyzed p21 protein levels. The knockdown of CD44-DT shows an upregulated p21 indicating that CD44-DT inhibits p21 protein degradation. Further, the indirectly impact of CD44-DT on p21 degradation analyzed by MG132 proteasome inhibition assay. In line with our hypothesis, in contrast to PCNA, p21 was markedly accumulated following siCD44-DT potently enhanced by treatment with MG132 (Fig. 5e). This suggests that the onco-lncRNA CD44-DT promotes the protein degradation of the p21 through PCNA stabilizing.

**Fig. 5).**
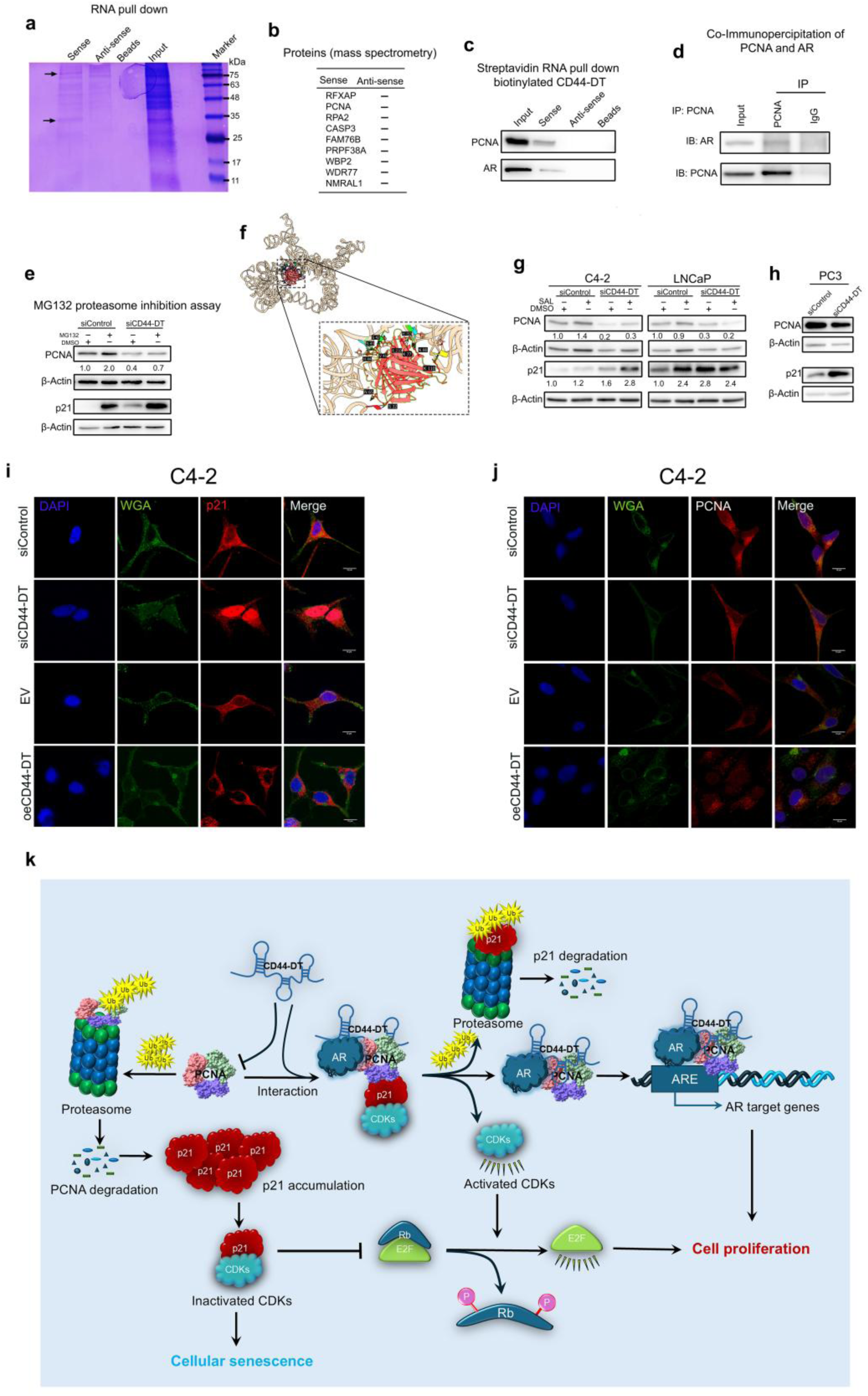
CD44-DT via interacting with PCNA-AR complex, regulates p21 function. a) Protein obtained from CD44-DT and its antisense RNA pull-down assays were analyzed in Coomassie blue-stained SDS-PAGE gel. b) The list of candidates CD44-DT interactive proteins that were existed in CD44-DT pull-down in comparison to the negative control conditions. c) Western blotting of PCNA and AR in protein retrieved from RNA pull-down assays. d) Co-IP reveals PCNA interacts with AR. e) Impacts of CD44-DT depletion on PCNA and p21 stability which was determined by MG132 treatment. f) HDOCK analysis of the interacting motifs between CD44-DT and PCNA. g) Western blotting analysis of the effect of CD44-DT KD on PCNA and p21 in SAL context. h) Western blotting analysis shows the effects of siCD44-DT on p21 in p53-deficent PC3 cells. i and j) Immunofluorescent staining of PCNA and p21 in C4-2 cell treated with siCD44-DT and oeCD44-DT. k) Schematic illustration shows that the mechanism of CD44-DT by which regulates the cellular senescence and cellular proliferation via interacting with PCNA-AR complex.

HDOCK analyses also predicted the potentially interaction between CD44-DT and PCNA (Fig. 5f). To further confirm the impact of CD44-DT on PCNA stabilization and p21 degradation, Western blotting experiments following knockdown and overexpression of CD44-DT in absence or presence of SAL were performed. Knockdown and overexpression experiments confirmed that CD44-DT promotes slightly PCNA protein stability but strongly inhibits p21 degradation, which might be the underlying molecular reasons to lead to cell proliferation (Fig. 5g, Fig.S5a). Interestingly, CD44-DT silencing in p53-deficient PC3 cells leads also to an increase in p21 protein levels and reduction in PCNA (Fig. 5h) suggesting that CD44-DT regulates the balance between p21 and PCNA in p53-independent manner, and explains the strong impact of CD44-DT on cancer cell growth and proliferation. Furthermore, to determine whether p21 has also feedback on PCNA, the impact of *CDKN1A* knockdown and overexpression on PCNA was analyzed. The data suggest that p21 does not impact on PCNA and there is no feedback loop between them (Fig.S5b). These findings suggest that CD44-DT by interacting with PCNA-AR-p21 complex acts as key regulators in cell decision between DNA replication or DNA repair through regulating PCNA and p21 at transcription and post-translational levels. This suggests that CD44-DT can be considered as a novel and critical lncRNA in PCa proliferation and a potential target for PCa treatment.

The correlation between CD44-DT and p21-PCNA complex might be originated from the subcellular localization of p21 and PCNA. Interestingly, it was suggested that p21 represents a dichotomous behavior in cells as it acts either as promoter or suppressor of cell cycle. In the nucleus, p21 suppresses proliferation and acts as a tumor suppressor, whereas its cytoplasmic localization promotes tumor proliferation and functions as an oncogene [40]. Strikingly, immunofluorescence analysis revealed a pronounced change in p21 subcellular localization in response to CD44-DT modulation. Whereas CD44-DT silencing results in nuclear p21 accumulation, the CD44-DT overexpression leads to a dramatically translocation of p21 into a cytoplasmic accumulation (Fig. 5i). Furthermore, PCNA subcellular localization was analyzed and IF data suggest that the localization of PCNA was also slightly changed but not as strongly as p21 in response to CD44-DT expression modulation (Fig. 5j). Collectively, our findings suggest the functional mechanistic of CD44-DT in stabilizing PCNA-AR complex and subsequently p21 degradation leads to cell proliferation (Fig. 5k)

### A novel chimeric lncRNA platform as an RNA-based PROTAC downregulates CD44-DT through AR degradation

Since AR is the key upstream factor of CD44-DT and also it interacts with CD44-DT-PCNA-p21 complex, has a critical role in PCa progression and targeting of this interaction has been implicated as a novel therapeutic strategy. Therefore, we rational designed a chimeric engineered ncRNA (cncRNA) as a novel chimeric ncRNA-based PROTAC to deplete AR protein in PCa cells and thereby further determine whether CD44-DT is regulated by AR. For that purpose, a cncRNA expression vector was constructed containing a fragment of HOTAIR, HOTAIR^948-1087^ and a fragment of SLNCR1, SLNCR1^597-637^ linked by a short linker. HOTAIR^948-1087^ contains a DZIP3 binding motif, whereas SLNCR1^597-637^ is responsible for binding to AR. DZIP3 is an RNA-binding E3 ubiquitin ligase that catalyzes the ubiquitination of substrate proteins, thereby targeting them for proteasomal degradation (Fig. 6a). This PROTAC cncRNA was designed to degrade AR by the proteasome.

**Fig. 6).**
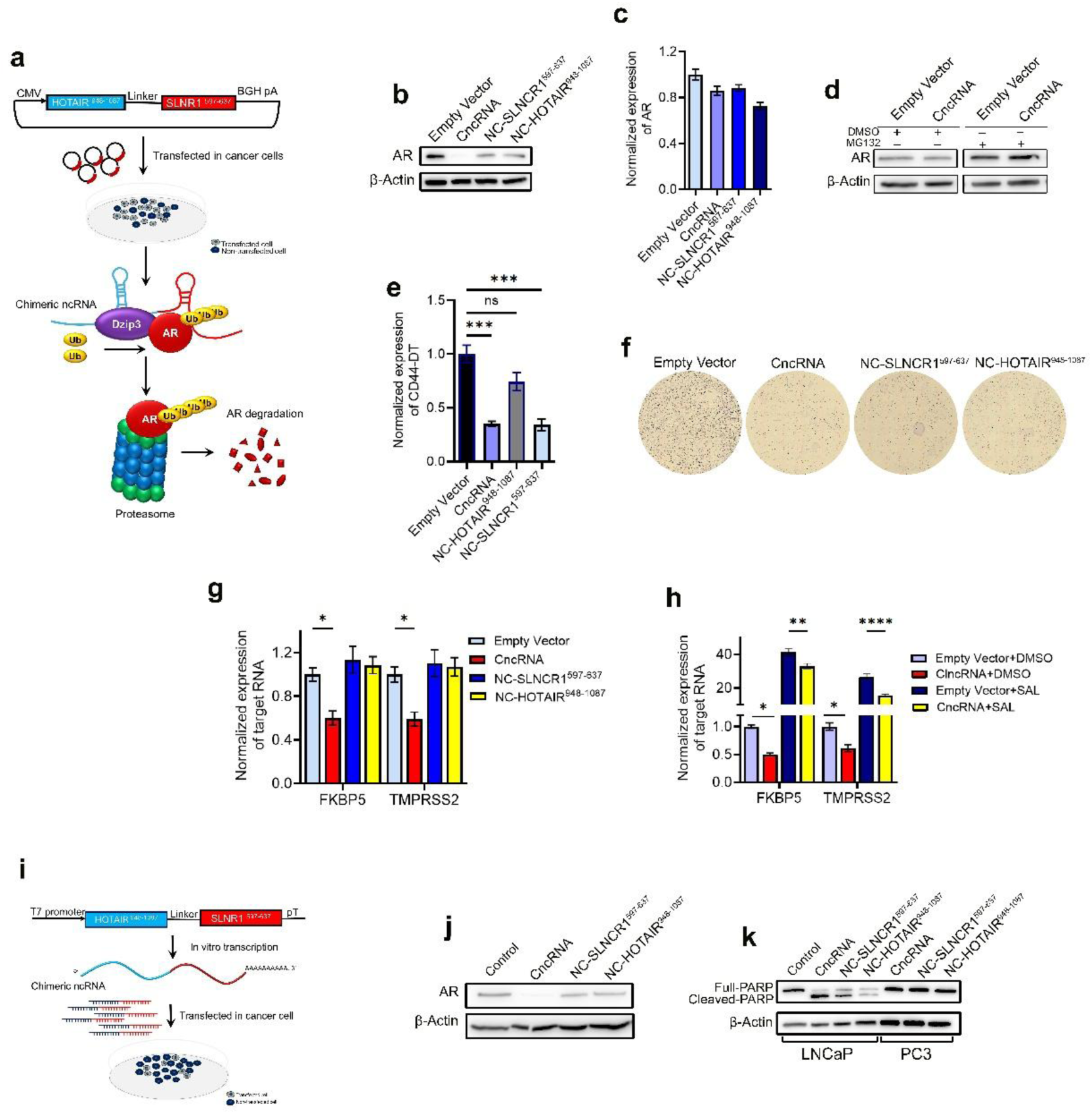
AR degradation by PROTAC chimeric ncRNA downregulates CD44-DT and induces apoptosis in PCa cells. a) Schematic illustration shows the design strategy of cncRNA. b) Analysis of AR expression at protein level by Western blot and c) at RNA level by qRT-PCR in C4-2 cells transfected with cncRNA and negative control vectors. d) Western blotting of AR in C4-2 cells transfected with cncRNA and control vectors following treatment with MG132. e) Bar graph shows the expression of CD44-DT in C4-2 cells when transfected with cncRNA and negative controls. f) Crystale violet staining represents the impact of cncRNA on C4-2 cells growth. g) Diagram shows the expression of AR target genes including FKBP5 and TMRPSS2 measured by qRT-PCR in C4-2 cells treated with cncRNA and negative control vectors. h) The expression of AR target genes was analyzed by qRT-PCR in C4-2 cells treated with cncRNA and negative control vectors in SAL context. Two-way ANOVA was performed for statistical analysis (*p ≤ 0.05, **p ≤ 0.01, ***p ≤ 0.001, and ****p ≤ 0.0001). i) Schematic illustration represents the design strategy for using chimeric RNA for transfection instead of vector-based overexpression. j) Confirming the reduction of AR expression at protein level after chimeric RNA transfection by Western blot. k) Analysis of PARP cleavage following cncRNA and negative controls transfection in LNCaP and PC3 cells.

Overexpression of this cncRNA by transfection with an expression vector resulted in significant reduction of AR at protein level without altering its RNA level in androgen-sensitive LNCaP cells (Fig. 6b and 6c). The ubiquitin-mediated degradation of AR was confirmed by MG132 treatment that leads to enhanced AR stability detected by Western blotting (Fig. 6d). In line with our hypothesis, CD44-DT was downregulated in cells transfected with cncRNA expressing vector suggesting CD44-DT expression is potentially regulated through AR probably being recruited to the enhancer within the first intron of CD44 in castration-sensitive cells (Fig. 6e). Consequently, LNCaP cell growth was also reduced. LNCaP cell transfected with cncRNA vector compared with cells transfected with the empty vector (Fig. 6f). Expression analysis showed that transcription of well-known AR target genes, *FKBP5* and *TMPRSS2*, are downregulated following cncRNA ectopic overexpression in LNCaP cells (Fig. 6g). Accordingly, cncRNA inhibited the expression of *FKBP5* and *TMPRSS2* in the presence of SAL (Fig. 6h). To improve the efficiency of transfection and better simulation the chimeric RNA-based treatment approach, the cncRNA was synthesized by in vitro transcription (Fig. 6i). RNA-transfection of the cncRNA resulted in reducing of AR at protein level (Fig. 6j). Interestingly, the rate of apoptosis strongly was increased compared to the negative controls. To decipher whether the apoptosis induction and growth inhibition effects of the cncRNA is due to AR degradation or not, we used the AR-negative PC3 cell line as a control. Our results showed that the apoptosis rate, determined by PRAP cleavage ratio, remained unchanged following the cncRNA transfection in PC3 cells (Fig. 6k). This suggests that the novel designed chimeric ncRNA PROTAC is specifically targeting the AR.

Together, disruption of AR signaling by PROTAC cncRNA leads to CD44-DT downregulation and consequently reduction of AR-positive PCa cell growth, introduces an innovative chimeric RNA platform that can be considered as a novel and creative targeted therapeutic strategy for PCa.

## Discussion

Despite the decline in PCa mortality due to improved early detection and advanced treatment, PCa remains still the second leading cause of cancer-related death in men in Western European countries [1, 41]. Dysregulation of androgen signaling plays a critical role in the initiation and progression of PCa and it appears the focus of androgen signaling has always been an important hormonal therapeutic target to treat PCa [27, 42]. Surprisingly, not only ADT but paradoxically also high dose androgen, SAL, restricts PCa growth by inducing senescence [10–12]. Recently, it has been shown that AR function is dependent on androgens dose, whereby low-dose androgens act through AR monomers and activate non-genomic mTOR signaling to promote proliferation, whereas high-dose androgens facilitate AR dimerization and consequently its translocation to the nucleus. The dimeric AR binds to androgen response elements (AREs) within the regulatory regions of target genes and recruits regulatory factors to activate or repress their transcription [27]. The majority of patients with PCa initially respond to ADT and AR antagonists, unfortunately most patients ultimately became resistant to these therapies and tumors progress into castration-resistant disease within several years, hence tumors grow despite low androgen levels [5]. The factors driving resistance to hormone therapy, and the exact mechanisms leading to evade hormone therapy, remain poorly understood. Our previous reports established that SAL-mediated cellular senescence as a tumor suppressive mechanism, has been regulated by signaling pathways such as p15^INK4b^/p16^INK4a^/pRb, AKT, and RBL2-DREAM complex [10–12, 24]. Recent reports showed that not only coding factors but also long non-coding transcripts contributed to therapy resistance and progression of PCa such as HOTAIR, PCGEM1 and MIR503HG [11, 20, 43].

Our analysis revealed that lncRNA CD44-DT expression is positively correlated with CD44 expression in most cancer types including PCa, likely due to its transcription from a shared BDP with CD44. Despite upregulation of CD44-DT in low-grade PCa, its expression showed a positive correlation with the expression of stemness-associated genes in PRAD. Interestingly, CD44-DT positively regulates the expression of core ESC-associated TFs including Oct4, Nanog, Sox2, and CD44 in PCa cells. However, previous studies reported that divergent transcripts (DTs) regulate the transcription of their protein-coding partner in pluripotent stem cells and they are appeared evolutionarily cooperate in the same signaling pathway [17, 19]. A recent report identified four subtypes of CRPC including AR-dependent (CRPC-AR), neuroendocrine (CRPC-NE), Wnt-dependent (CRPC-Wnt) and stem cell-like (CRPC-SCL) subtypes. CRPC-SCL subtype expressed high-level of CD44 and driven by the AP-1 complex [14]. We suggested that SAL-induced CD44-DT, enables unexpectedly some cells to escape senescence, promote tumor plasticity and progress to CRPC-SCL which can identified by upregulation of CD44-DT and ESC-associated TFs.

Transcriptome analysis showed that CD44 was upregulated in patients who received BAT. In line with this result, CD44-DT was also upregulated in SAL-treated PCa cells and C4-2 xenograft mice models treated with SAL, identifying CD44-DT as a novel androgen-responsive lncRNA. PRAD dataset analysis further revealed that CD44-DT, as a novel AR direct target gene, correlates with hallmark pathways promoting PCa growth, tumor progression and therapy resistance including NF-κB, p53, PI3K-AKT-mTOR, and androgen response signaling pathways. Consistently, RNA-seq analysis also showed that the expression of CD44-DT is associated with the hallmark pathways and interestingly this correlation is enhanced in SAL context. These findings introduced CD44-DT as a novel target of AR that plays critical role in tumor progression and therapy resistance.

In this study, CD44-DT identified as a potent suppressor of SAL-induced senescence that is upregulated following SAL treatment. In contrast to SAL, CD44-DT markedly suppresses cellular senescence as verified by both KD and overexpression experiments in 2D adherent and 3D spheroid models. Consistently, transcriptome analysis showed that the pathways repressed by siCD44-DT are enriched in oncogenic programs comparable to those repressed by SAL. RNA-seq analysis also showed that CD44-DT triggers critical therapy-resistant cellular plasticity pathways leading to PCa progression such as Wnt, AP-1, Hedgehog, NFĸB and p53 signaling. Thus, we promoted to investigate the mechanistic function of CD44-DT in reduction of SAL-induced cellular senescence in PCa cells.

Our previous reports established that pRb-E2F1 and non-genomic AR signaling PI3K/AKT mediate SAL-induced senescence [11, 12]. We found that SAL-induced CD44-DT downregulates p21 and p27, key cell cycle inhibitors, induces Rb phosphorylation, and consequently removes inhibition of E2F1, leading to cell cycle progression and suppression of SAL-induced cellular senescence. Accordingly, our data showed that CD44-DT confers resistance to SAL-induced senescence through suppressing AKT phosphorylation and that of its downstream target, p70S6K. Thus, the upregulation of CD44-DT in response to SAL suggests that this upregulation triggers PCa progression through enabling cancer cells to escape senescence which may result in cellular plasticity and cancer cell dedifferentiation.

Deciphering the upstream regulatory mechanism of androgen-dependent CD44-DT upregulation appears to be critical for understanding how androgen signaling upregulates CD44-DT that leads to reduction of SAL-induced senescence. This regulatory mechanism is important because enhancing SAL-induced senescence has been considered as a promising therapeutic strategy. Accordingly, androgen ATAC-seq data suggest that AR may reprogram a tightly packed, inaccessible chromatin region into a loose, open state at a regulatory region within the first intron of CD44. Consistent with the ATAC-seq data analysis, ChIP-seq results showed that AR is recruited at this genomic region and consequently activates transcription of CD44-DT. Motif analysis showed an AR binding motif nearby Fox binding motif in this region, likely contributed in activation of CD44-DT transcription.

Previous studies showed that AP-1 complex is a known upstream activator of CD44 [15, 35] and it differentially acts in different CRPC subtypes due to the differential expression patterns of its members in a context-dependent manner [14]. Accordingly, FOS and FOSL2, two members of AP-1 family, along with CD44-DT are upregulated following SAL treatment. Consistently, ATAC-seq and ChIP-seq data confirmed that the AR recruitment at cis-regulatory elements of these members of AP-1 genes which led to their upregulation following SAL. We initially hypothesized that AP-1 complex activates transcription of CD44-DT in PCa cells. Interestingly, while AP-1 inhibition by T5224 treatment notably downregulated CD44-DT in LNCaP and C4-2 cell lines, yet had no effect on CD44-DT expression in PC3 cells, which showed that high-level expression of CD44-DT. Although, CD44-DT promoter chromatin appears to be more accessible in PC3 cells, it isn’t regulated by AP-1, and T5224-sensitivity of PC3 is not related to this promoter activation. In contrast, downregulation of CD44-DT in C4-2 and LNCaP cells treated with T5224 suggesting AP-1 regulates the expression of CD44-DT in cooperating with AR signaling in AR-positive PCa cells. Taken together, the AR/AP-1 axis may act as upstream regulator of CD44-DT that leads to its upregulation. Therefore, complete disruption of the androgen signaling axis is still an important therapeutic target to treat PCa.

To decipher the exact mechanism of CD44-DT, RNA pull down assay revealed that CD44-DT directly interacts with PCNA and AR. Previous reports have shown that PCNA mediates p21 ubiquitylation and degradation to regulates the balance between DNA replication and DNA repair [38, 39, 44]. CD44-DT interacts with and stabilizes PCNA-AR complex leads to its accumulation in nucleus. Accumulated PCNA also binds to p21 and promotes it to ubiquitination and degradation results in Cdk activation and subsequently Rb phosphorylation that leads to E2F releases and activates the transcription of cell cycle progression genes. AR-PCNA complex also induces proliferation via upregulating AR target genes. Strikingly, CD44-DT silencing strongly accumulated p21 in the nucleus and its overexpression dramatically depleted almost of p21 from nucleus of cancer cells suggest CD44-DT may act as a key oncogene in opposition of p53, a well-known tumor suppressor gene. The similar impacts of CD44-DT suppression on p21 expression in p53-defienct PC3 cells suggests CD44-DT can be functioned in p53-independent manner and it can potentially regulate the p21 expression in p53-independent pathway. Our findings suggest that CD44-DT acts as a pivotal onco-lncRNA responsible for homeostasis p21 levels via PCNA stabilizing in p53-indepenet manner. Subsequently CD44-DT may control PCNA-AR complex to regulate the cell fate decision between DNA repair or DNA replication which leads to cancer cell proliferation or cell cycle arrest-induced cellular senescence. Therefore, this complex that scaffolds with CD44-DT can be considered as a target for PCa target therapy.

Recently, lncRNAs are noticed particularly promising therapeutic tools because of their capability to interact with various biomolecules in the cell including proteins, chromatin, mRNA and miRNAs. RNA molecules have been considered as a novel therapeutic targets or tools aimed to combat different diseases including neurodegenerative diseases, viral infections and cancer [21, 45].

In the early 1980s, Humulin, as a first recombinant therapeutic protein, was generated by DNA recombination technology for diabetes therapy [46, 47]. During these years, chimeric fusion proteins such as romiplostim and alefacept has also been approved by FDA for clinical use, treats immune thrombocytopenia (ITP) and plaque psoriasis, respectively [48, 49]. Therefore, we here designed an innovative chimeric ncRNA (cncRNA) that able to sponge the AR protein and promote its ubiquitination, thereby directing it to the proteasome for degradation. This PROTAC cncRNA has been constructed from functional protein interaction domains derived from HOTAIR and SLNCR1 that was successfully able to ubiquitinate and degrade AR proteins leading to a dramatic reduction in CD44-DT expression. Surprisingly, cancer cell proliferation was incredibly reduced and apoptosis rate was dramatically increased following AR degradation by the cncRNA, introducing a new creative strategy for developing novel RNA therapeutic approaches for PCa treatment. For many years, chimeric and recombinant therapeutic proteins have been widely used as therapeutic agents for the treatment of various diseases, including cancer [50]. Here, we introduce an innovative creative strategy for engineering chimeric ncRNAs, establishing a novel class of RNA therapeutics for cancer treatment. This approach expands the therapeutic potential of RNA-based medicines and provides a new avenue for specifically targeting pathogenic proteins and their regulatory pathways in different diseases including cancer.

In summary, our findings identify CD44-DT as a novel AR target gene that represses SAL-induced senescence and potentially acts as a crucial early driver of the growth of AR-positive PCa through AR/AP-1 axis. CD44-DT suppression through various approaches including siRNA-mediated knockdown, AP-1 inhibitor (T5224) and clncRNA dramatically enhances SAL-induced cellular senescence and reduces the growth of AR-positive PCa cells. These results introduce CD44-DT as a novel onco-lncRNA that drives PCa progression, and also as a novel potentially therapeutic target for the development of targeted PCa treatment.

## Authors’ contribution

M.H.A and A.B. wrote the draft and revised it. M.H.A and A.B. designed and supervised the study. F.H. and M.H.A performed experiments and data collection. G.A.R. performed the animal experiments. K.S. performed slicing of tumor spheroids. F.H. performed bioinformatic analysis. J.K., F.H. and M.H.A. analyzed the ATAC-seq and ChIP-seq data. N.R. performed mass spectroscopy and analyzed that. All the authors read and approved the submitted version.

## Data availability

The RNA-seq, ATAC-seq and ChIP-seq datasets, and PDX xenografts analyzed in this study are accessible in GEO under accession numbers GSE229555, GSE247593 and GSE232554 respectively. The TCGA PRAD and GTEx prostate datasets from GEPIA and Starbase were used for correlation analysis (assessed on 06 Jun 2025).

## Consent for publication

Not applicable.

## Data Availability

Data required to assess the decisions of this research are available in the main text or the supplementary tables and figures. The RNA-seq data generated in this study have been deposited in the GEO database.

## Acknowledgments

The author gratefully acknowledges the Alexander von Humboldt Foundation for support this research. We thank Dr. Emilio Cirri and the Core Facility Proteomics at the Leibniz Institute on Aging-Fritz Lipmann Institute (FLI), Jena, Germany for excellent Mass spectroscopy support.

