## Supplementary for "CD44-DT regulates p21-dependent senescence via interacting with AR-PCNA complex, revealing a therapeutic vulnerability to chimeric ncRNA-mediated AR degradation in prostate cancer"

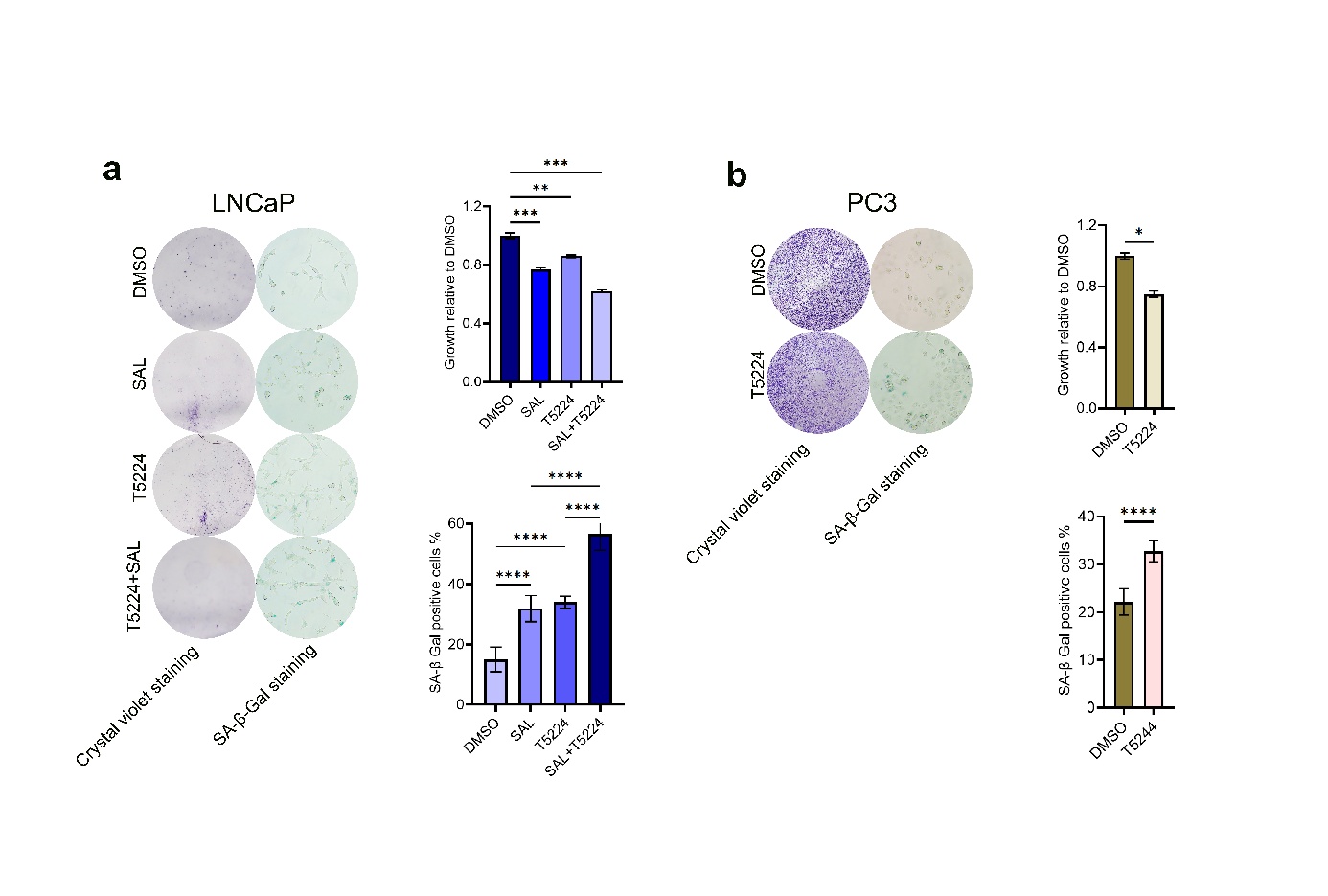

**Fig. S1) AP-1 inhibitor, T5224, reduces the growth and induces the cellular senescence in PCa cells. a)** Crystal violet and SA-β-Gal staining shows the impact of T5224 and SAL on growth and senescence of castration-sensitive, androgen-dependent LNCaP cells and **b)** AR-negative PC3 cells.

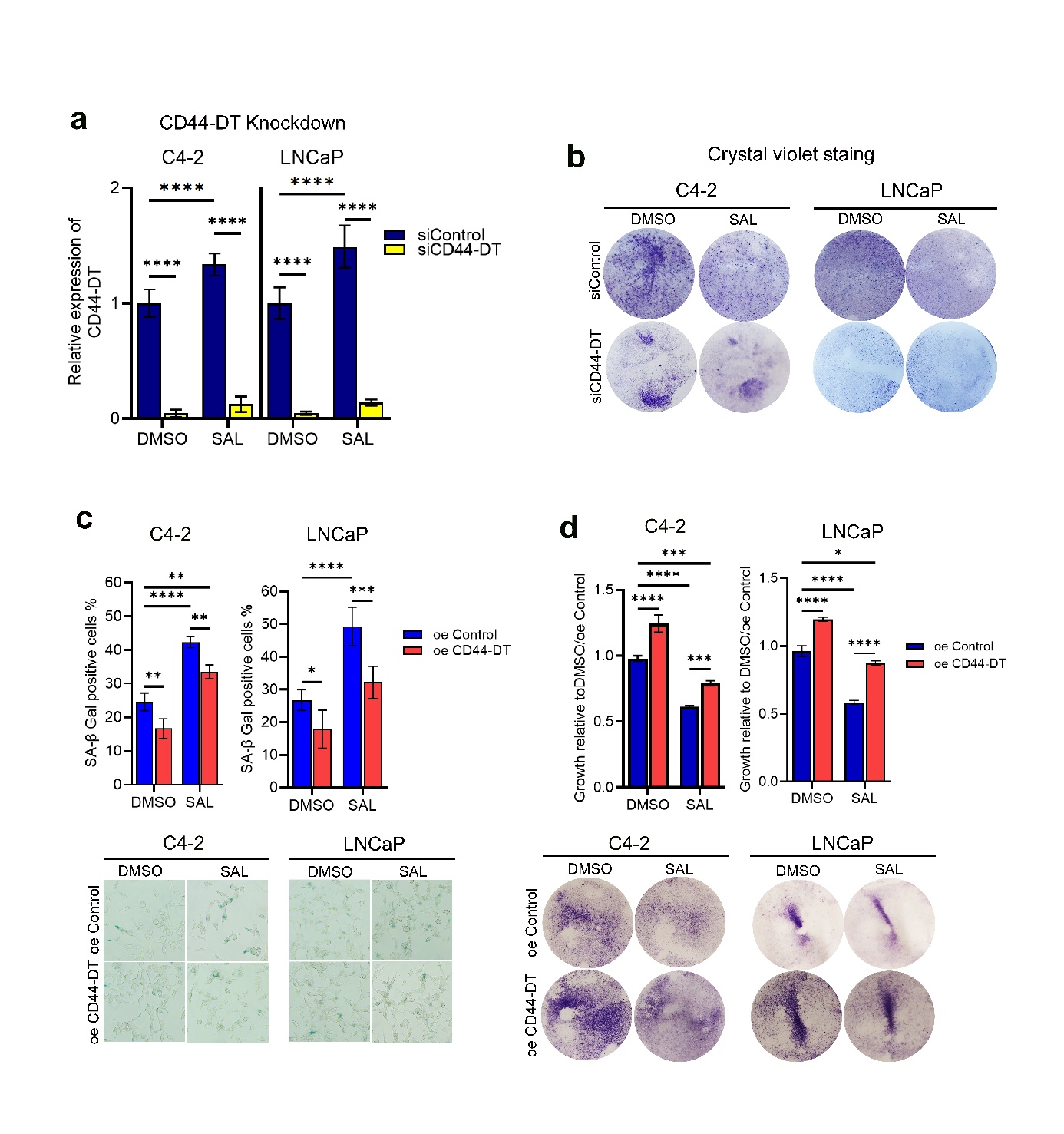

**Fig. S2) CD44-DT represses cellular senescence and induces PCa cells growth.** **a)** CD44-DT expression was analyzed using qRT-PCR in CD44-DT KD cells relative to scrambled siRNA transfected cells in the presence of SAL. Data represent the mean ± SEM. **b)** SA-β-Gal staining of siRNAs transfected C4-2 and LNCaP cells treated with SAL or DMSO. **c)** The the percentage of senescent cells of PCa cells determined by SA-β-Gal staining ollowing oeCD44-DT transfection and SAL treatment. **d)** Analysis of growth ratio when CD44-DT was ectopic overexpressed in C4-2 and LNCaP cells in presence of SAL by crystal violet staining. Two-way ANOVA was performed for statistical analysis (*p ≤ 0.05, **p ≤ 0.01, ***p ≤ 0.001, and ****p ≤ 0.0001).

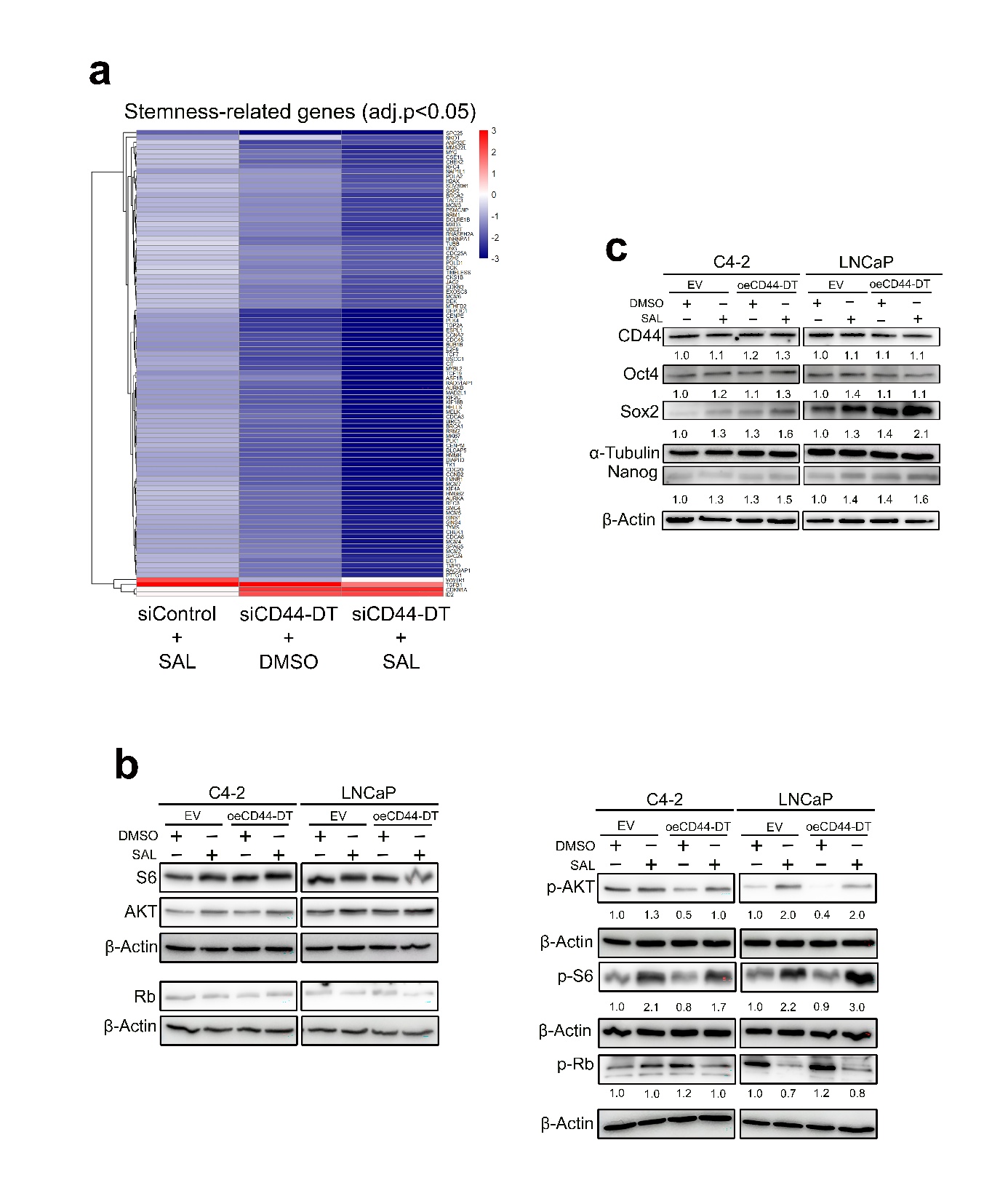

**Fig. S3) The impacts of CD44-DT on PCa growth via regulating stemness-associated genes and PI3K/AKT/mTOR and pRb-E2F1 signaling pathways. a)** Heatmap shows siCD44-DT downregulates stemness-associated genes more profoundly compared to SAL. **b)** Analysis of core ESC-associated TFs at protein level by Western blot following CD44-DT overexpression in presence of SAL. **c)** Western blot shows that CD44-DT overexpression modulates the phosphorylation of AKT, S6 and Rb in C4-2 and LNCaP cells. Number represents the relative band intensity that was normalized to β-Actin (n = 3).

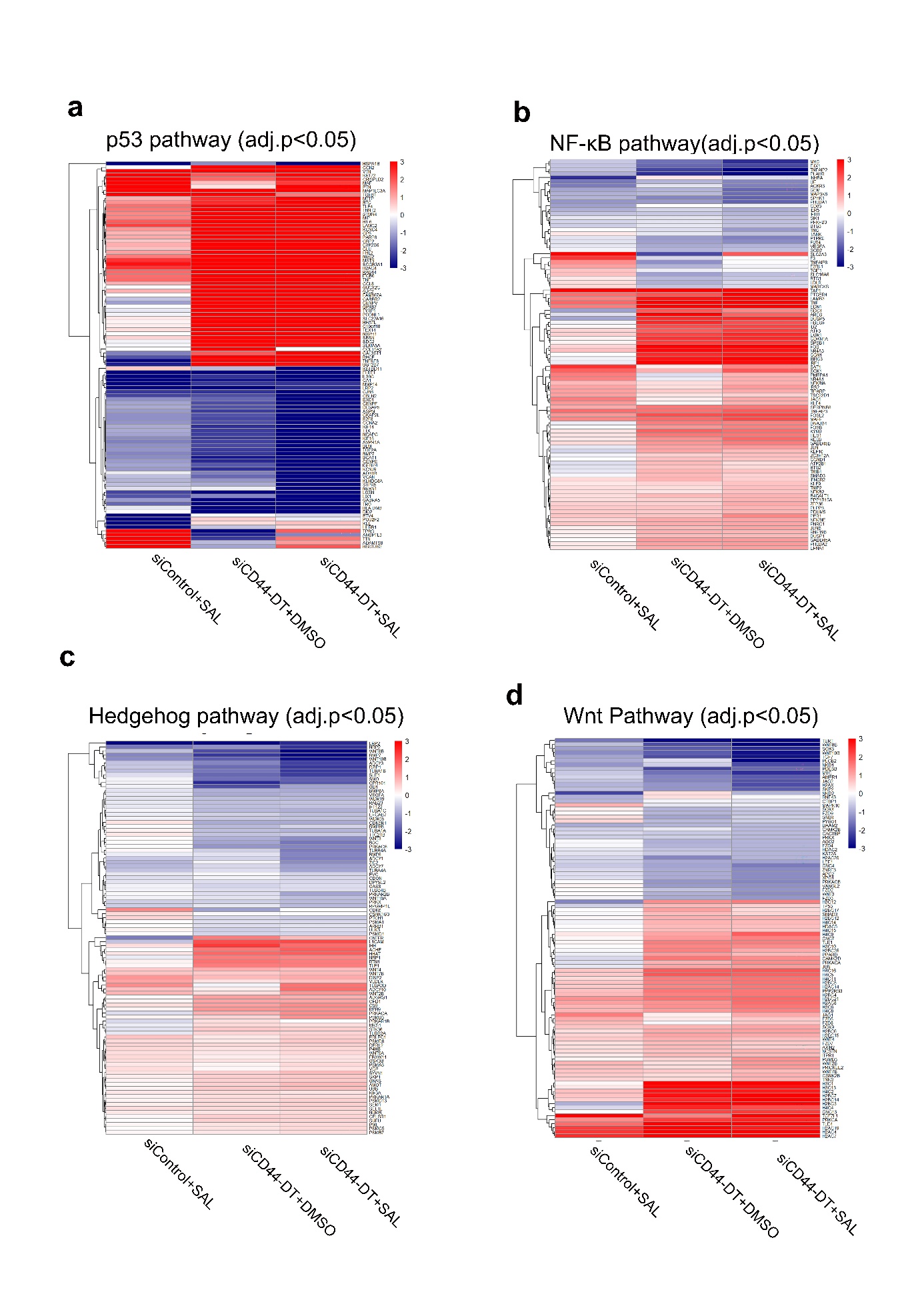

**Fig. S4) CD44-DT acts as key factor in tumor plasticity and progression of PCa. a)** Heatmaps representing the critical role of CD44-DT in regulating the PCa tumor plasticity and progression through P53, **b)** NF-ĸB, **c)** Hedgehog and **d)** Wnt pathways. Heatmaps were extracted from RNA-seq data generating on the Nanostring nCounter platform in SAL-treated C4-2 with or without siCD44-DT versus cells treated with DMSO plus siControl.

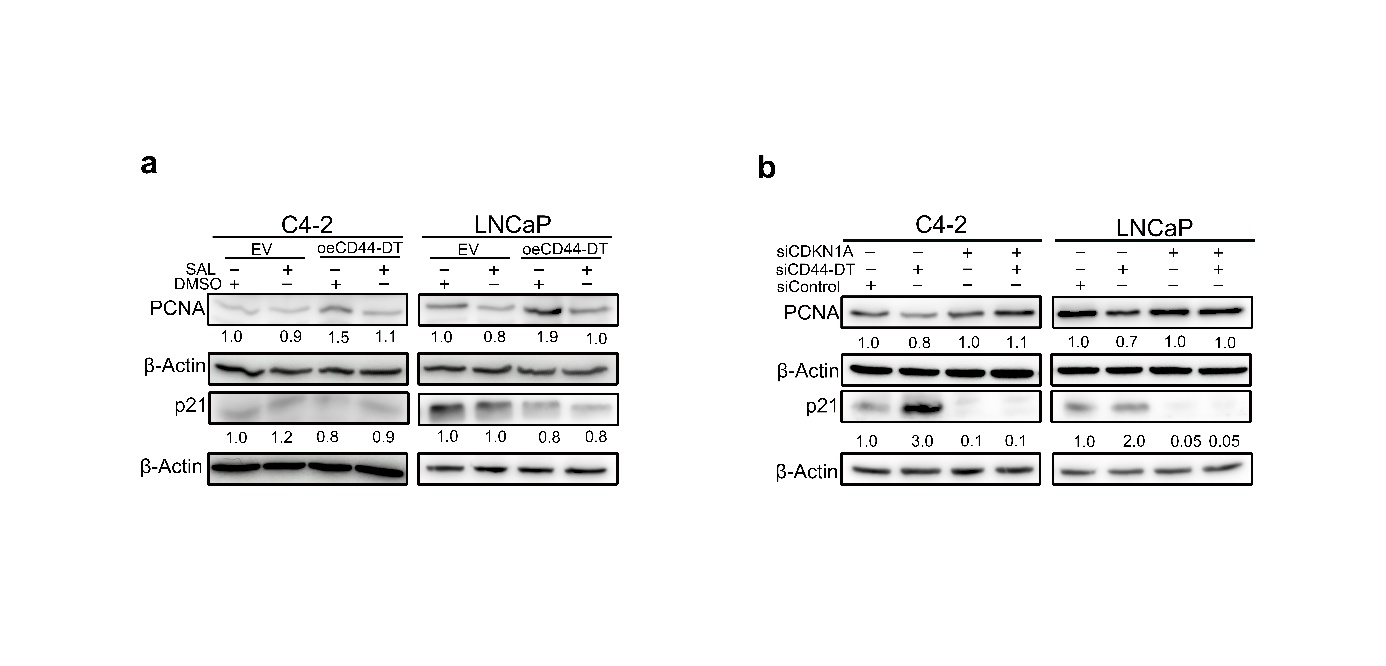

**Fig. S5) CD44-DT/PCNA/AR complex acts as an upstream regulator of p21. a)** CD44-DT overexpression upregulated PCNA and consequently reduced p21 but **b)** PCNA abundance remained unchanged after CDKN1A knockdown in PCa cells.
